# IL-1-driven stromal-neutrophil interaction in deep ulcers defines a pathotype of therapy non-responsive inflammatory bowel disease

**DOI:** 10.1101/2021.02.05.429804

**Authors:** Matthias Friedrich, Mathilde Pohin, Matthew A. Jackson, Ilya Korsunsky, Samuel Bullers, Kevin Rue-Albrecht, Zoe Christoforidou, Dharshan Sathananthan, Rahul Ravindran, Raphael Sanches Peres, Hannah Sharpe, Kevin Wei, Gerald F. M. Watts, Elizabeth H. Mann, Alessandra Geremia, Tom Thomas, Moustafa Attar, Oxford IBD Cohort Investigators, Roche Fibroblast Network Consortium, Sarah McCuaig, Lloyd Thomas, Elena Collantes, Holm H. Uhlig, Stephen Sansom, Alistair Easton, Soumya Raychaudhuri, Simon P. Travis, Fiona M. Powrie

## Abstract

Current inflammatory bowel disease (IBD) therapies are ineffective in a high proportion of patients. Combining bulk and single-cell transcriptomics, quantitative histopathology, and *in situ* localisation, we describe heterogeneity of the tissular inflammatory response in IBD treatment failure. Among inflammatory pathotypes, we found high neutrophil infiltration, activation of fibroblasts, and vascular remodelling at sites of deep ulceration was a feature of non-response to several anti-inflammatory therapies. Activated fibroblasts in the ulcer bed display neutrophil chemoattractant properties that are IL-1R- but not TNF-dependent. The identification of distinct, localised, tissular pathotypes associated with treatment non-response will aid precision targeting of current therapeutics and provide a biological rationale for IL-1 signalling blockade in ulcerating disease.

## Introduction

Inflammatory bowel diseases (IBDs) are a heterogeneous group of disorders characterised by inflammation throughout the gastrointestinal tract. The aetiology involves maladaptation between the host and its intestinal microbiota, a dialogue controlled by genetic and environmental factors ^1^. The complex multi-factorial nature of IBD is partly reflected in its clinical phenotypes, encompassing both Crohn’s disease (CD) and ulcerative colitis (UC), and in a range of microscopic features such as granulomas, lymphoid aggregates, crypt abscesses, and ulcers ^2,3^. Treatments for IBD include general immunosuppressants (such as corticosteroids), immunomodulators (such as thiopurines), or biologics that target specific inflammatory mediators ^4^. Amongst the latter, tumor necrosis factor α (TNF) targeting is most common ^5^ but alternate approaches, such as blockade of leukocyte homing to the gut (anti-integrin α4β7 (vedolizumab)) are increasingly used ^6^. Although anti-TNF therapy has revolutionised the treatment of IBD, identifying the patients who will respond remains a major challenge. Up to 40% of IBD patients are primary non-responders and for a substantial fraction of initial responders, treatment will later lose efficacy ^7-9^.

Our previous work identified high expression of the IL-6 family member Oncostatin M (OSM), and its receptor OSMR, in the inflamed intestine of IBD patients as associated with non-response to anti-TNF therapy ^10^. Notably, OSM produced by leukocytes signals primarily into stromal cells such as fibroblasts and endothelial cells. Subsequent bulk and single-cell transcriptomic studies have associated cell subsets of fibroblasts, inflammatory mononuclear phagocytes (MNP), neutrophils, and pathogenic T- and plasma cells with therapy non-response in both UC and CD ^11-17^. However, these studies have not identified the processes underlying quantitative changes in cellular ecology nor how they affect treatment response. It is also not known whether the cellular and molecular determinants of treatment response are uniform across non-responsive patients or if several different tissular pathologies promote therapy failure through distinct mechanisms. Similarly, little is known about the mechanistic determinants of therapy response that are shared/unique to individual drugs. Further understanding in these areas is crucial to design personalised treatment regimens and new therapeutics for individuals that do not respond to current options.

In this study we dissected the cellular and molecular landscape of inflamed tissue in IBD patients by integrating whole-tissue and single-cell gene expression profiling with quantitative *in situ* analyses and functional *ex vivo* assays. We then explored how individual signatures of tissue inflammation associate with non-response to specific treatments. Transcriptomic changes were found to reflect changes in the tissular response and characterised by distinct histological features. Most notably we identified a pathotype in a subset of patients that was associated with non-response across several current IBD therapeutics. Tissues from those patients are characterised by a high abundance of tissue neutrophils, the activation of a neutrophil-attractant program in fibroblasts, (peri-) vascular cell expansion, and enhanced IL-1 signalling at sites of deep ulceration. These functional definitions of disease provide a basis for rational targeting of existing medications and a novel mechanistic avenue to target inflammation in non-responsive patients displaying ulceration with fibroblast and neutrophil remodelling.

## Results

### Identification of gene co-expression signatures of cellular ecology in inflamed IBD tissue

The surgical removal of inflamed tissue becomes a therapeutic option in IBD when medical therapies have failed. Using such tissues as our discovery cohort, we examined gene expression profiles in difficult-to-treat IBD. From the 31 IBD patients (n=8 UC, n=22 CD and n=1 IBDu) from which samples were collected, only 20% were treatment-naïve (i.e. had a resection as primary therapy for localised disease), while 48% had received two or more different medications before the time of surgery (Supplementary Table 1). Amongst the n=41 tissue samples from these patients, n=15 were classified as (macroscopically) uninflamed, including n=7 samples for which paired uninflamed/inflamed tissue was available. Weadditionally used unaffected, non-tumour tissue collected from colorectal cancer patients undergoing surgery as non-IBD controls for comparison (n=39). Bulk RNA sequencing (RNAseq) was used to generate whole tissue gene expression profiles across all samples (n=41 IBD and n=39 non-IBD; ‘discovery cohort’).

To identify sets of genes reflective of distinct biological processes, we applied weighted gene correlation network analysis (WGCNA) to cluster co-expressed genes in an unbiased manner across all tissue samples. This identified 38 modules of highly co-expressed genes (M1-M38) (Supplementary Table 2). We correlated the expression (module eigengene) of these modules with sample characteristics, clinical phenotypes and histologic (microscopic) inflammation (Nancy Index ^18^); 28 modules were significantly associated with at least one of these measures (Figure 1A). Modules were found to have dichotomous associations with traits. About half of the modules had significant positive correlations with histologic inflammation, whilst the others had significant negative associations (Figure 1A). Fewer and less strong correlations were observed between module expression levels and other metadata, such as the intestinal sampling site or IBD subtype (CD or UC), and overall these measures displayed correlations in a similar direction to histologic inflammation. Age appeared to have similar associations as inflammation but this was an artefact of the older nature of the non-IBD samples used as controls. In a paired analyses of only inflamed and uninflamed IBD tissue samples from the same patients (n=7), the difference in expression of a module between tissue pairs remained highly correlated with the module’s association with histologic inflammation (Nancy Index) (R=0.8, P<0.001, Extended Data Figure 1A), confirming that these co-expression modules reflected inflammatory processes.

**Figure 1.**
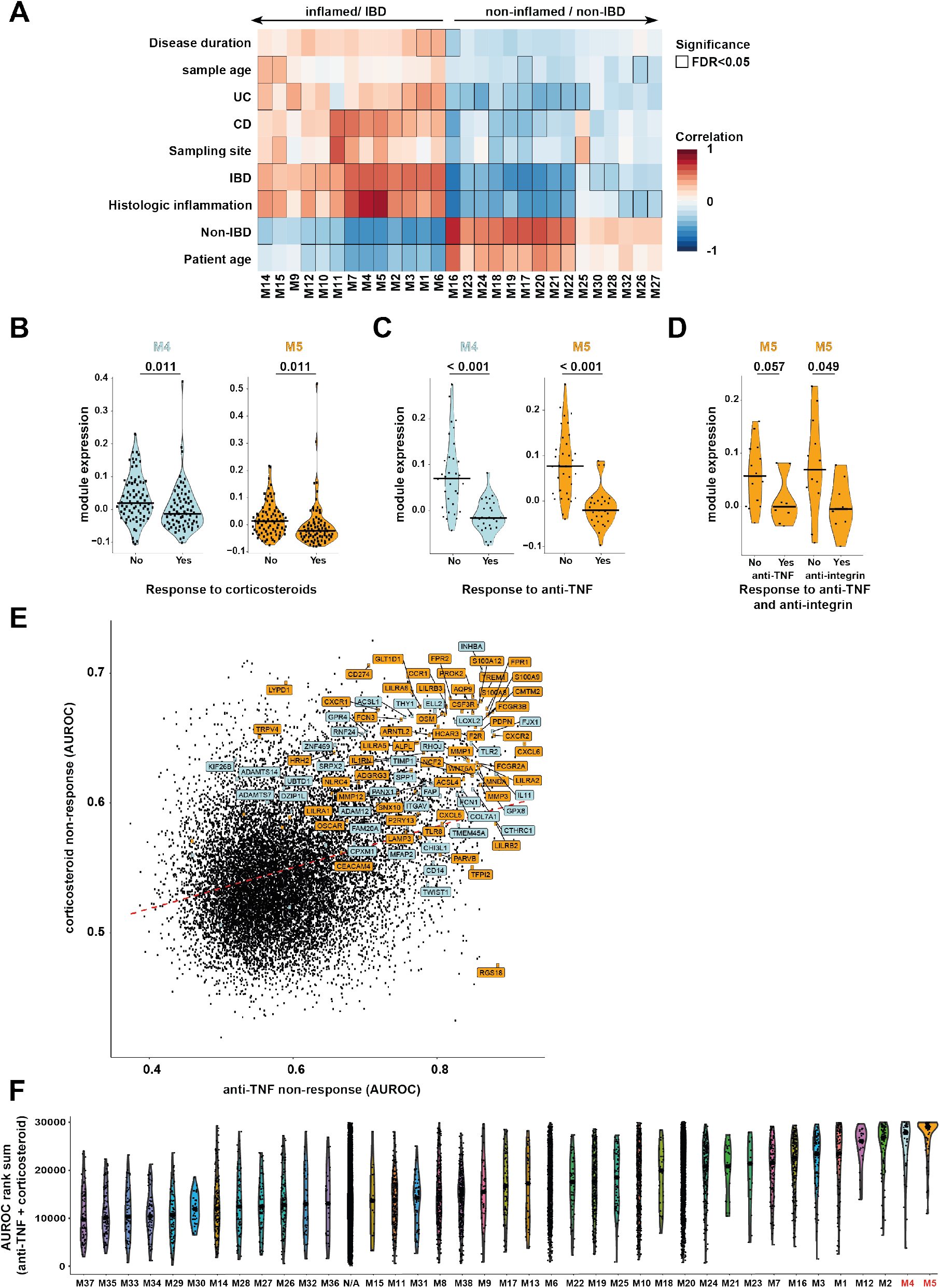
Identification of gene co-expression signatures of inflammation associated with patient non-response to multiple different IBD therapies. A) Pearson correlation between module eigengenes and clinical and histologic metadata in inflamed and CRC derived tissues within the discovery cohort; all modules/features with at least one significant association shown, bordered squares indicate significant correlations (FDR p<0.05). B-D) Module M4 and M5 expression (eigengene value) in non-responders and responders before the start of corticosteroid (B ^15^), anti-TNF (C+D ^20,21^) or monthly anti-integrin therapy (D ^21^) (Mann-Whitney U, FDR adjusted P-values, post-hoc to ANOVA comparisons across the various treatment regimens in the Arjis 2018 study in the case of D). E) Performance (AUROC) of individual genes for predicting non-response to corticosteroid (y-axis) and anti-TNF (x-axis) therapy; genes contained in M4 and M5 are labelled and highlighted by turquoise and orange datapoints respectively. F) Violin plots showing the rank of genes based on their predictive power (area-under-the-receiver-operator-curve, AUROC) for response to both anti-TNF and corticosteroid therapy, comparing all modules as detected in the WGCNA analysis. Combined ranks represent the sum of each gene’s ranks in the separate corticosteroid and anti-TNF analyses (their ranks on the x and y axes in (E)).

To determine whether the gene co-expression patterns we detected reflected changes in the cell type composition of patient tissues, we applied *in silico* cell type deconvolution analysis to the RNAseq data of our discovery cohort (*xCell*, ^19^). Correlating predicted cell type scores with module expression (eigengenes) (Extended Data Figure 1B), modules positively correlated with histologic inflammation (Figure 1A) were associated with signatures of stromal cells (e.g., fibroblasts), mononuclear phagocytes (e.g., M2 macrophages), B-lymphocytes and plasma cells, T-lymphocytes (e.g., CD8+ T-cells), and granulocytes (e.g., neutrophils). Modules negatively correlated with histologic inflammation were predicted to reflect epithelial cells, smooth muscle cells and M1 macrophages. These results suggest that the co-expression patterns that we observed to be associated with inflammation were, at least in part, being driven by differences in the cellular composition of the inflamed tissues.

### Co-expression signatures of inflammation predict patient response to IBD treatments

Given previous associations between the expression of individual genes and cell types with therapy response in IBD, we aimed to determine if our inflammation-associated gene co-expression signatures represented biological features relevant to treatment outcomes. We projected all of the modules onto whole tissue gene expression data derived from prospective studies of response to anti-TNF, corticosteroid, or anti-integrin therapy ^15,20,21^. At least 79% of the genes within each module could be identified in the three datasets, enabling accurate quantification of the modules within them (Extended Data Figure 1C and Supplementary Table 3). The expression of 15 modules was significantly (adjusted p<0.05) higher in non-responders to anti-TNF prior to treatment (n=61 total patients in the study). Seven modules were significantly higher in non-responders (and one significantly lower) in the corticosteroid study (n=206 patients) and two modules were higher in non-response to anti-integrin therapy (n=20 patients) (Supplementary Table 4).

Strikingly, across all three therapy-response datasets, each involving different therapeutics, modules M4 and M5 were consistently amongst the strongest associations with non-response in pre-treatment samples (Supplementary Table 4 and Figures 1B, 1C and 1D). This overall trend of increased expression in non-responders was significant in meta-analyses of both M4 (p=0.0025, standardised mean difference (SMD)=0.87, 95%CI=0.31-1.44) and M5 (p=0.0123, SMD = 0.88, 95%CI=0.19-1.58) across the different treatments.

To determine if the associations with non-response are uniform across the genes in modules M4 and M5 or driven by a small number of highly predictive genes, we compared the ability of all genes individually to predict response to anti-TNF and corticosteroid therapy. This again revealed that genes from modules M4 and M5 were amongst the top predictors of non-response to both anti-TNF and corticosteroid therapy relative to those in other modules (Figure 1E, Figure 1F, Supplementary Table 5). Thus, M4 and M5 reflect a coordinated shift in the expression of all their constituent genes in relation to therapy non-response. Overall, M4 and M5 were consistently the top predictors of non-response across multiple IBD medications.

### Co-expression modules linked with therapy non-response represent distinct histopathologic features

As well as predicting therapy response, modules M4 and M5 also demonstrated the strongest correlation with histologic inflammation in the discovery cohort, as defined by the Nancy score ^18^ (Figure 1A). Using an additional clinical cohort of Oxford UC patients (Supplementary Table 6), we confirmed that the Nancy score is higher in non-responders to anti-TNF therapy before the start of treatment (Figure 2A). Interestingly, this was not true for the UCEIS (an endoscopic score of mucosal inflammation) or other clinical or endoscopic measures (Extended Data Figure 2A).

**Figure 2.**
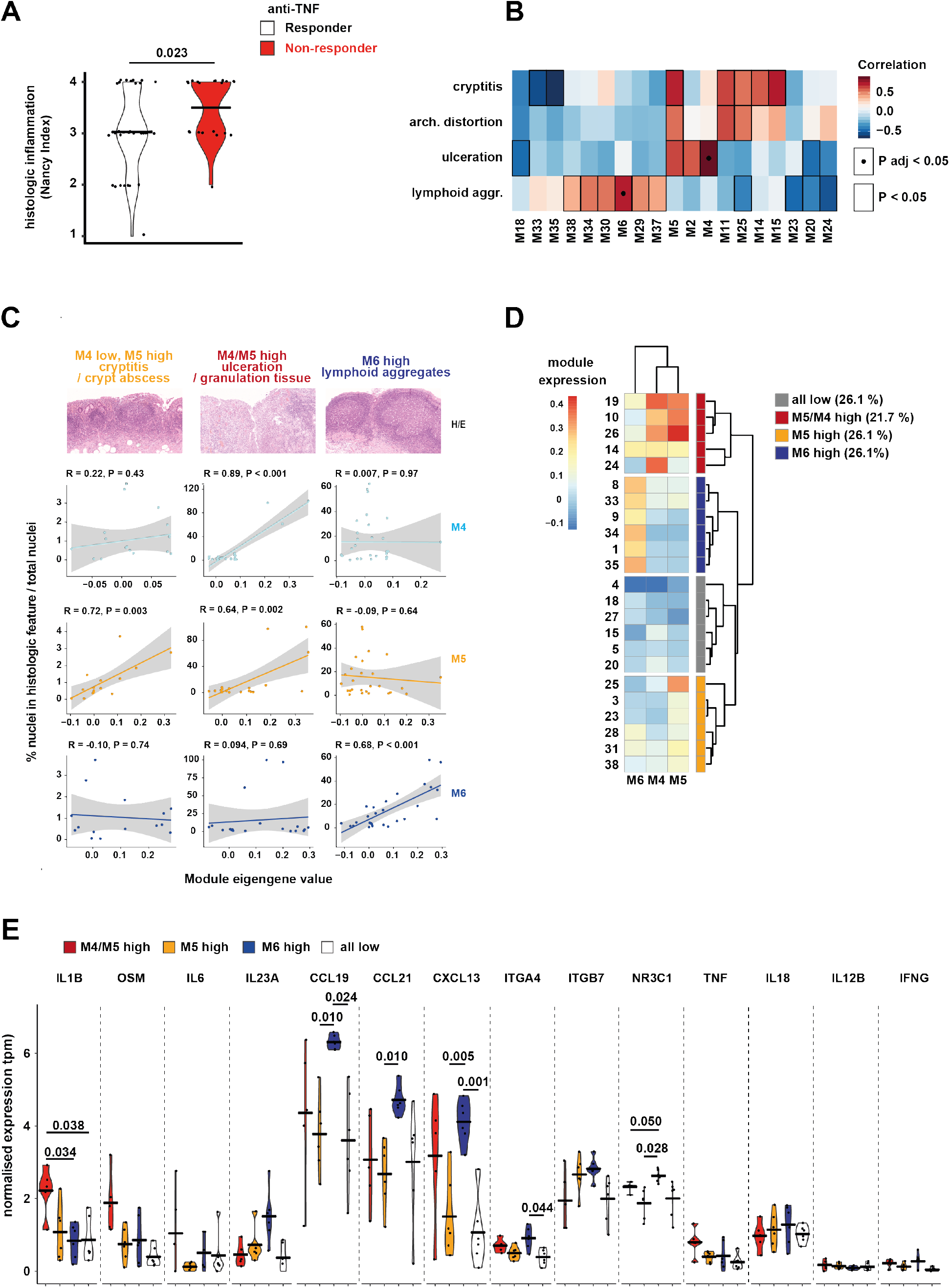
Co-expression modules linked with therapy non-response represent distinct histopathologic features. A) Nancy histologic scores in non-responders to anti-TNF therapy before the start of treatment (horizontal bars indicate geometric mean, Mann-Whitney U test P values given). B) Heatmap of correlations between module eigengene expression and histological features quantified across tissues from IBD patients in the discovery cohort. Nominally significant associations (p<0.05) are indicated by borders and FDR significant (FDR p<0.05) associations are indicated by dots. C) Scatter plots showing eigengene expression of M4, M5 and M6 versus selected quantified histologic features in tissue samples from IBD patients of the discovery cohort. D) Classification of M4/M5 high, M5 only high, M6 high and M4/M5/M6 low patients in the discovery cohort, based on hierarchical clustering of module eigengene values from inflamed tissue samples. E) Normalised expression (tpm) of cytokine and therapeutic target genes that were reliably (in >50% of samples) detected in the discovery cohort. The expression of these genes is compared in the M4/M5 high (red), M5 only high (orange), M6 high (blue) or M4/M5/M6 low tissues (bottom panel). Horizontal lines indicate the median and p-values (Wilcoxon signed rank test, adjusted for multiple testing) for each comparison are given if significant (P < 0.05).

On this basis we postulated that the gene co-expression patterns in the dataset, which we previously linked to changes in cellular composition, might also reflect the manifestation of histopathologic features in patient tissues. To explore this, we quantified established histopathologic features of IBD on H&E sections of resected patient tissue (Extended Data Figure 2B). First, we examined the correlation of histopathologic features with each other (Extended Data Figure 2C). The only strong positive correlations observed were between cryptitis/crypt abscess and architectural distortion/ goblet cell depletion; as well as several associations with granulomas, although the latter estimates were based on very few cases where both granulomas and the other features were observed. We then looked for correlations between the expression of the co-expression modules and histologic features scored from tissue where both were available (n=36). Several nominally significant associations were observed between modules and various features (Figure 2B); however, only positive correlations between M4/ulceration and M6/lymphoid aggregates remained significant after adjusting for multiple testing (P adjusted < 0.05, Figure 5A). Notably, the relation of these two inflammation-associated modules was almost orthogonal, each correlating only with one of the features (Figure 2C). Despite not reaching significance after correction, M5 – also highly correlated with the Nancy score (Figure 1A) – correlated strongly with both ulceration and cryptitis/crypt abscesses (Figure 2C). We also confirmed the associations of M4, M5 with ulceration in an independent paediatric cohort (n=172) containing inflamed tissues of both UC and CD patients {Haberman, 2014 #722; Loberman-Nachum, 2019 #1093) (Extended Data Figure 2D). In this dataset, 11% of all patients with IBD showed high M4/M5 tissue expression (Extended Data Figure 2E). Similar to our dataset, M6 expression was not significantly different by ulceration status in the paediatric cohort, although we noted that overall M6 expression was also much lower in these biopsy samples (see discussion).

The almost orthogonal relation of M4/M5/ulceration with M6/lymphoid aggregates suggested these may represent distinct underlying inflammatory processes that may be more or less dominant in a given patient’s tissue. To investigate this, we grouped patients by unsupervised clustering on module M4/M5/M6 expression to determine the relative proportion of samples belonging to these groups. This yielded four groups: M4/M5 high expression (21.7% of patients), M6 high expression, M5 only high expression and M4/M5/M6 low (each 26.1% of patients) (Figure 2D). We then plotted the expression of cytokines and therapeutic targets reliably detected in our discovery cohort across these groups (Figure 2E). The M4/M5 high group displayed significantly increased expression of *IL1B* compared to the rest of the patients (Figure 2E, red). However, neither *ITGA4*/*ITGB7* (targeted by anti-integrin), *N3RC1* (targeted by corticosteroids) nor *TNF* (targeted by anti-TNF) were increased in the tissue of these patients (Figure 2E). By contrast, high expression of module M6 was linked to increased levels of *ITGA4* and *N3RC1*, as well as *CCL19, CCL21* and *CXCL13* but not *TNF* (Figure 2E, blue).Patients high in M5 expression only did not demonstrate significant changes in cytokine/therapeutic target signatures (Figure 2E, orange). These results suggest that patient responses to specific treatments might be determined by which inflammatory pathology predominates at the tissue level. M4/M5-high tissues did not show any increase in current therapeutic targets and these modules were associated with non-response to all therapies tested; whereas M6-high tissues only showed no increase of *TNF* expression in the tissue, which corroborated our previous association analyses where it was only associated with non-response to anti-TNF (Supplementary Table 4).

The quantification of histological features confirmed that an increased expression of both M4 and M5 is linked to the presence of deep ulcerations and M5 to cryptitis/crypt abscesses. Whereas other inflammatory features were instead correlated with alternative co-expression patterns, as in the case of lymphoid aggregates and the M6 module. Patients with ulceration and high M4/M5 expression showed no significant up-regulation of genes targeted by the current medications, but an increase in *IL1B*, warranting further exploration of the mechanisms underlying this signature.

### High expression of modules M4 and M5 reflects neutrophil infiltrates, activated fibroblasts and epithelial cell loss

We performed a more detailed exploration of the changes in cellular composition and activation state that produce the M4 and M5 co-expression module signature. Our *in silico* cell type deconvolution revealed that M4 and M5 were predominantly associated with stromal cells, such as fibroblasts, and granulocytes, such as neutrophils (Extended Data Figure 1B). We confirmed this by projecting modules M4 and M5 onto single-cell transcriptomic datasets derived from inflamed and non-inflamed CD {Martin, 2019 #567} and UC patient tissue ^12^. This showed that the module M4 likely reflected the presence of “activated/inflammatory fibroblasts”, whereas module M5 reflected “myeloid cells/inflammatory monocytes” (Figures 3A and 3B).

**Figure 3.**
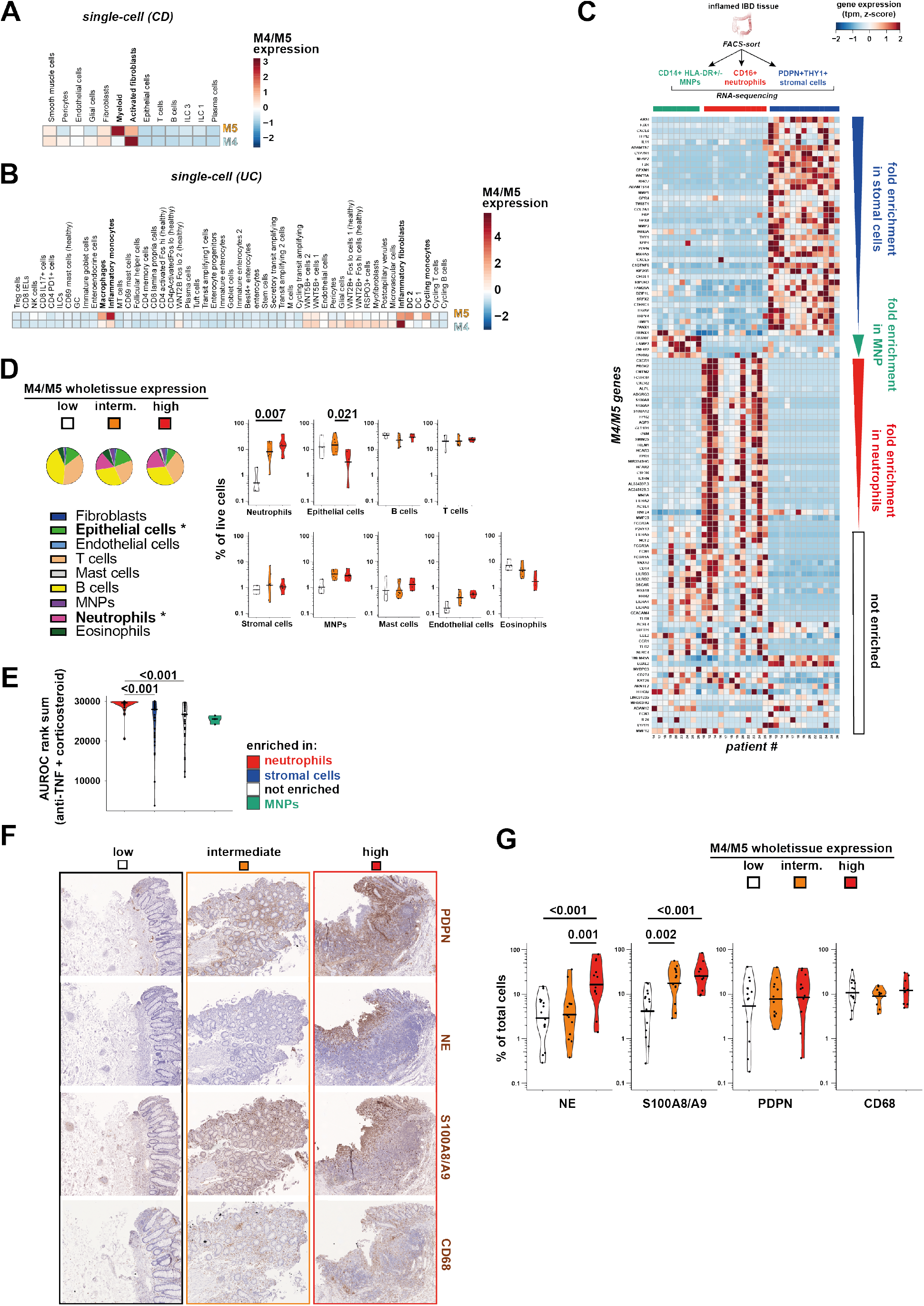
High expression of modules M4 and M5 reflects neutrophil infiltrates, activated fibroblasts and epithelial cell loss. A+B) Module M4 and M5 expression in cell clusters detected by scRNAseq in UC ^12^ and CD ^11^ patient tissue. C) Heatmap of the expression (TPM values, z-score, Manhattan distance clustering) levels of all genes contained within M4 and M5 in THY1+PDPN+ stromal cells, CD16^hi^ neutrophils, and CD14+HLA-DR+/- MNPs, FACS-sorted from inflamed IBD patient tissue. The genes are ordered by their log fold-change of significant enrichment (P adjusted < 0.05) in either cell type. D) FACS cell type percentages in tissue isolates from IBD patients, classified into low (white), intermediate (orange) or high (red) expression of M4/M5 (see Extended Data Figure 3E). Pie-charts show medians across samples and boxplots individual samples. (* in pie charts indicates cell population percentages significantly different between groups, post-hoc 2-way ANOVA adjusted P-values are given, if significant). E) Violin plots showing the rank of genes based on their predictive power (AUROC) for response to both anti-TNF and corticosteroid therapy, comparing genes significantly enriched in neutrophils, stromal cells, MNPs or neither. Combined ranks represent the sum of each genes ranks in the separate corticosteroid and anti-TNF analyses. F) Illustrative IHC staining (DAB, counterstain hematoxylin) of Podoplanin (PDPN), neutrophil-elastase (NE), calprotectin (S100A8/A9) or CD68 in serial sections of IBD patient tissue classified as low, intermediate or high for M4/M5 whole tissue gene expression (see Extended Data Figure 3F). G) Automated quantification (% positively stained cells of total cells detected in inflamed areas) of IHC stainings as shown in F); each staining was performed on inflamed tissue sections with low (n=17), intermediate (n=13) and high (n=12) M4/M5 whole tissue gene expression (see Extended Data Figure 3F); post-hoc 2-way ANOVA adjusted P-values are given, where significant.

Given that cell type deconvolution correlated neutrophil scores with M5, but the single-cell datasets (which did not capture neutrophils) correlated monocytes/macrophages with M5 genes, we aimed to identify genes within M4 and M5 enriched in either of these cell types, as well as in stromal cells. We FACS-sorted well-defined hematopoietic and non-hematopoietic cell subsets from the intestinal tissue of non-IBD and IBD patients and measured the expression of selected M4/M5 genes by qPCR (see Extended Data Figure 3A for gating strategy). Several M4/M5 genes were highly expressed in CD16^hi^ neutrophils and PDPN+THY1+ stromal cells (Extended Data Figure 3B). This was confirmed by targeted RNAseq from neutrophils, stromal cells and mononuclear phagocytes (MNP) (see Extended Data Figure 3C for gating strategy), which were bulk-sorted from inflamed endoscopic biopsies of n=13 IBD patients (UC and CD, Figure 3C, see Supplementary Table 7 for patient cohort details). We first carried out pathway analysis to assign function to all the genes (including those not in M4 and M5) enriched in either cell type (see Supplementary Table 8 for differential gene expression analysis). As expected, this demonstrated that neutrophils were enriched in anti-microbial and tissue-toxic granule biology when compared to MNPs that were mostly defined by genes belonging to the antigen presentation pathway (Extended Data Figure 3D). Stromal cells were enriched in many genes assigned to extracellular matrix pathways (Extended Data Figure 3D). Of all differentially expressed genes between the cell types, n=39, n=31 and n=4 of all genes contained in M4/M5 (n=110) were significantly enriched in stromal cells, neutrophils and MNPs, respectively (Supplementary Table 8, Figure 3C). Compared to both neutrophils and MNPs, sorted stromal cells were enriched in transcripts for components (*COL7A1*) and remodelling enzymes (*MMP1/3, ADAMTS7/14*) of the extracellular matrix (ECM), markers of activated fibroblasts (*THY1, PDPN, FAP*), as well as for ligands of the chemokine receptors CXCR1/CXCR2 identified as enriched in neutrophils (*CXCL5/6*). Genes encoding major neutrophil chemokine receptors (*CXCR1/2)*, subunits of the antimicrobial peptide calprotectin (*S100A8/A9)*, receptors for IgG immunoglobulin constant regions (*FCGR3B)* and the cytokine *OSM*, which we previously linked to non-response in IBD ^10^, were enriched in neutrophils. Amongst the four genes enriched in MNPs, *CD300E* is a marker of activated monocytes ^22^, whereas *LAMP3* has been described as indicative of mature dendritic cells ^23^. The enrichment of many M4 and M5 genes in sorted neutrophils explained the high correlation of modules M4 and M5 with the Nancy index (Figure 1A), which is weighted by the abundance of neutrophils for scoring ^18^.

Since differences in whole tissue gene expression signatures could be driven by both changes in the transcriptional profiles within cell-types and/or overall changes in cell-type composition, we used flow cytometry to test whether the number of neutrophils and fibroblasts correlated with M4 and M5 tissue expression (see Extended Data Figures 3E for classification of tissues by M4/M5 expression). The percentage of neutrophils was significantly increased (up to 10 fold) in M4/M5 high tissues while the percentage of stromal cells remained unchanged (Figure 3D). Additionally, epithelial cells were significantly decreased in M4/M5 high tissues (Figure 3D). We also observed non-significant trends for an increase of MNP and endothelial cells, as well as a trend for decreased eosinophils with high M4/M5 expression (Figure 3D). Whilst these trends may become significant with an increased number of samples, we noted that neutrophils accounted for up to 38% of the total live cells in the M4/M5 intermediate and high group, whilst the percentage of MNPs was much lower (<5%) (Figure 3D). Additionally, we found that the M4/M5 genes significantly enriched in neutrophils and stromal cells, but not MNPs, demonstrated highest predictive power for non-response to anti-TNF and corticosteroids (Figure 3E).

FACS counts can be biased by tissue digestion methods, so we also quantified the presence of neutrophils, stromal cells and MNPS *in situ* by immunohistochemical staining of resected formalin-fixed, paraffin-embedded (FFPE) inflamed tissue from IBD patients (Figure 3F). Again, IBD tissues with high expression of M4 and M5 in whole tissue (see Extended Data Figure 3F for classification) demonstrated a higher percentage of Neutrophil Elastase-(NE) and Calprotectin-(S100A8/A9) positive cells, but not PDPN-positive stromal cells or CD68+ MNPs, in inflamed tissues (Figure 3G). This further confirmed that M4/M5 high tissue harbours an increased number of neutrophils which stain positive for NE and S100A8/A9. High M4/M5 expression in the whole tissue thus reflects ulceration characterised by a dominance of neutrophil infiltration, expression of genes characteristic of activated fibroblasts, and the loss of epithelial cells.

### M4/M5 gene expression is associated with neutrophil-attracting fibroblasts and endothelial and perivascular cell expansion

M4/M5 high patients are characterised by a high abundance of neutrophils but the modules also contain many genes indicative of activated stromal cells (Figure 3C). Furthermore, whilst the number of PDPN+ stromal cells was not increased, we observed an increased overall staining intensity of PDPN in M4/M5 high patients (Figure 3F). We therefore hypothesised that the stromal signatures in M4 and M5 arise from altered activation states (including the upregulation of PDPN) and/or changes of cellular composition within the stromal compartment that correlate with the infiltration of neutrophils. To dissect this relationship, we applied single-cell sequencing to EPCAM-CD45-intestinal stromal cells from endoscopic biopsies of inflamed UC patients (n=7) and healthy donors (n=4) (see supplementary Table 7 for patient cohort details), and compared tissues with low, intermediate and high M4/M5 expression (Figure 4A+B and Extended Data Figure 4A). As expected, tissues from all healthy donors were M4/M5 low, as well as the tissue from one IBD patient with a low histological inflammation score (Nancy score=1). We used Harmony to integrate all single-cell datasets and account for inter-patient and inter-sequencing batch effects ^24^. Overall, 6 stromal clusters were obtained, which we assigned to endothelial cells (*ACKR1*+*CD34*+), pericytes (*NOTCH3*+*MCAM*+), myofibroblasts (*MYH11*^*hi*^*ACTG2*+) and three clusters of fibroblasts: *PDGFRA*^*high*^*PDPN*^*low*^*SOX6*+ (“PDGFRA+”) fibroblasts, *PDGFRA*^*low*^*PDPN*^*low*^*ABCA8*+ (“ABCA8+”) fibroblasts and *CD90*^*hi*^*PDPN*^*hi*^*PDGFRA*+*ABCA8*+*FAP*+ “inflammatory” fibroblasts, based on the top differentially expressed markers and previously described annotations ^11,12,25^ (Figure 4C and Supplementary Table 9). *PDPN* was expressed by myofibroblasts and all three fibroblast clusters, with highest expression found in inflammatory fibroblasts. *THY1* (CD90) was highly expressed in pericytes and inflammatory fibroblasts and expressed at lower levels in ABCA8+ fibroblasts (Figure 4C).

**Figure 4.**
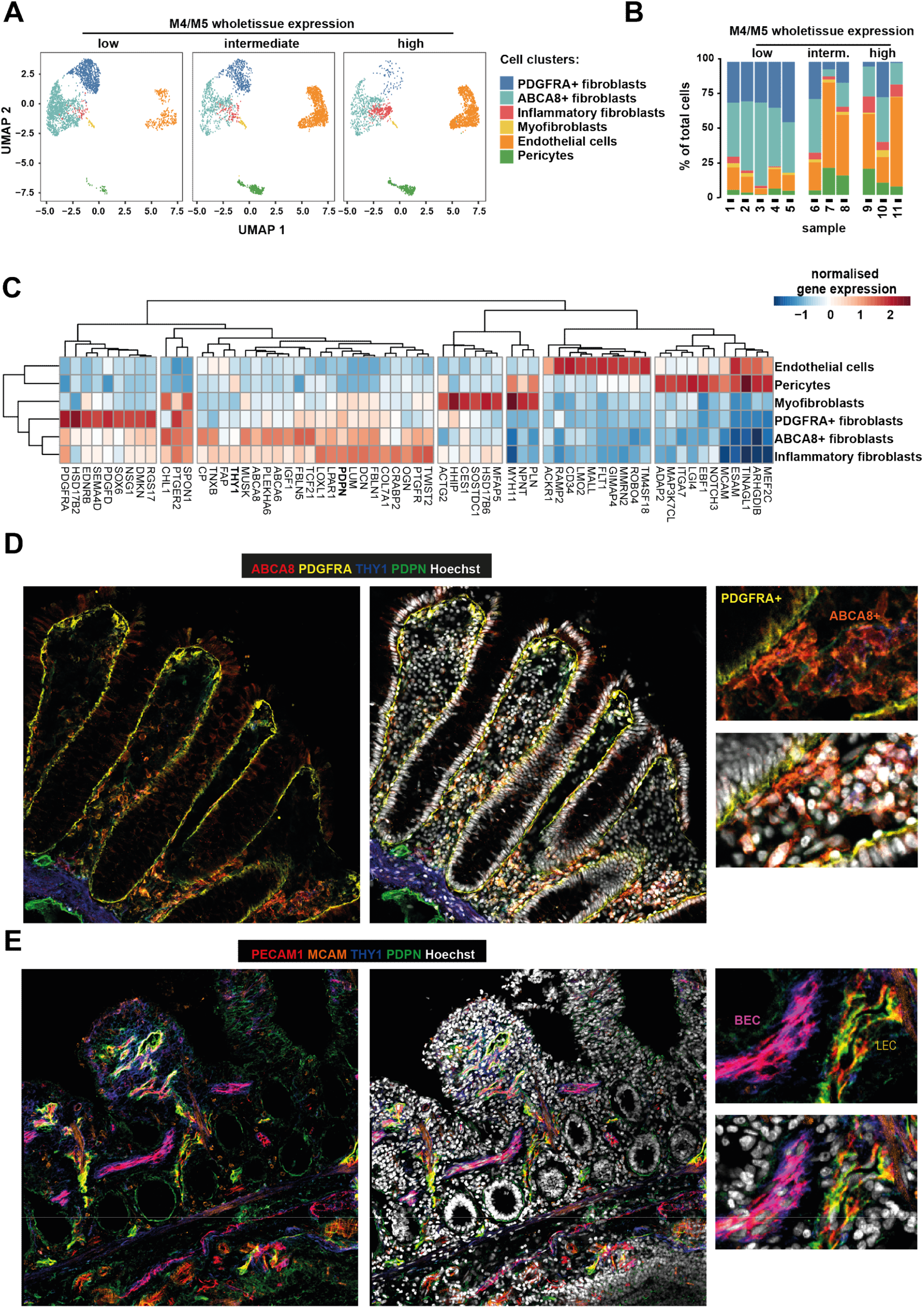
Stromal architecture of the large and small intestine in health and disease. A) UMAP of stromal clusters identified by Harmony in stromal compartments FACS-sorted from healthy donor and IBD patient tissue with low, intermediate and high M4/M5 whole tissue gene expression (see Extended Data Figure 4A). B) Proportion (% of total stromal cells) of the cell type clusters in A in the M4/M5 low, intermediate and high tissue. C) Heatmap of selected markers of each of the cellular cluster as in A, as identified by Harmony; Expression values are normalised log2 fold changes (Wald statistic 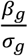 from DESeq2 analyses. D) Immunofluorescent staining of THY1 (Blue), Podoplanin (PDPN, Green), ABCA8 (red) and PDGFRA (yellow) to visualise the localisation of fibroblast subsets in resected tissue from IBD patients (uninflamed areas). E) Immunofluorescent staining of THY1 (Blue), Podoplanin (PDPN, Green), PECAM1 (Red) and MCAM (orange) to visualise the localisation of vascular (endothelial) and perivascular cells (uninflamed areas). PDGFRA: PDGFRA+ fibroblasts, ABCA8+: ABCA8+ Fibroblasts, BEC, blood endothelial cells, LEC: lymphatic endothelial cells.

Next, we developed a panel of antibodies for *in situ* analysis of intestinal tissue to confirm that the clusters of stromal cells detected by transcriptomics represented spatially separated cell subsets. The panel comprised anti-THY1, anti-PDPN, anti-PDGFRA and anti-ABCA8 to localise the different subsets of fibroblasts (Figure 4D and Extended Data Figure 4B). Anti-THY1 and anti-MCAM were used to localise pericytes, and anti-PECAM1 (CD31) to localise endothelial cells (Figure 4E and Extended Data Figure 4C). In uninflamed large (colon) and small (ileum) intestinal tissue, high PDGFRA staining was observed in sub-epithelial fibroblasts, which also stained low for PDPN (Figure 4D and Extended Data Figure 4B). By contrast, ABCA8 stained a distinct fibroblast population residing in the intestinal lamina propria (Figure 4D and Extended Data Figure 4B). Highest PDPN staining was found on lymphatic endothelial cells (PECAM1+ and PDPN+, Figure 3E and Extended Data Figure 4C). By contrast, THY1 formed a gradient of staining intensity from the perivascular niche toward the lamina propria (as recently described in ^26^), being expressed by both ABCA8+fibroblasts and MCAM+ pericytes, as well as cells in the muscular layer of the submucosa (Figures 3D,E and Extended Data Figure 4B, C).

We then characterised how the identified stromal compartments differed across IBD patient tissues with either low, intermediate or high M4/M5 expression. In the single-cell dataset, the percentage of inflammatory fibroblasts, pericytes and endothelial cells was increased in the M4/M5 intermediate and high patient groups at the expense of ABCA8+ and PDGFRA+ fibroblasts (Figures 4A+B, Extended Data Figure 4D). FACS analysis verified that PECAM1+ endothelial cell and PDPN+FAP+ inflammatory fibroblast frequencies were increased within the stromal compartment in inflamed tissue, compared to uninflamed adjacent tissue (Extended Data Figure 4E).

To see which of those clusters contributed most to M4/M5 expression, we projected the genes contained in the M4/M5 modules onto our scRNAseq data. Notably, the highest expression of M4 was detected in the inflammatory fibroblast cluster, suggesting the emergence of this cell cluster as an underlying process in M4/M5 high IBD patient tissue (Figure 5A). Within M4/M5 high tissues, neutrophil-targeting CXCR1/CXCR2 ligands CXCL1, CXCL2, CXCL3, CXCL5, CXCL6 and CXCL8 were significantly higher in inflammatory fibroblasts in comparison to other clusters (Figure 4B and Supplementary Table 10). We also identified several genes indicating extracellular matrix remodelling (MMP1, MMP3, MMP13) and previously identified markers associated with inflammatory fibroblasts (IL11, IL24, FAP) as higher in inflammatory fibroblasts compared to other stromal cells (Supplementary Table 10). Within the cluster of inflammatory fibroblasts, *PDPN, FAP, CXCL1, CXCL2, CXCL3, CXCL5, CXCL6* and *CXCL8* were also significantly increased in M4/M5 high compared to M4/M5 low and intermediate tissues, whereas *ABCA8* expression was downregulated (Supplementary Table 11). Nevertheless, ABCA8 fibroblasts and PDGFRA fibroblasts both still expressed the above-mentioned chemokines in the M4/M5 intermediate and high groups (Figure 5B), raising the possibility that the inflammatory fibroblast cluster represents an activation state of ABCA8 fibroblasts or/and PDGFRA fibroblasts. Indeed, trajectory (pseudotime) analysis indicated that inflammatory fibroblasts may represent a transcriptomic state in between ABCA8+ and PDGFRA+ fibroblasts (Extended Data Figure 4F) and could potentially arise from either population.

**Figure 5.**
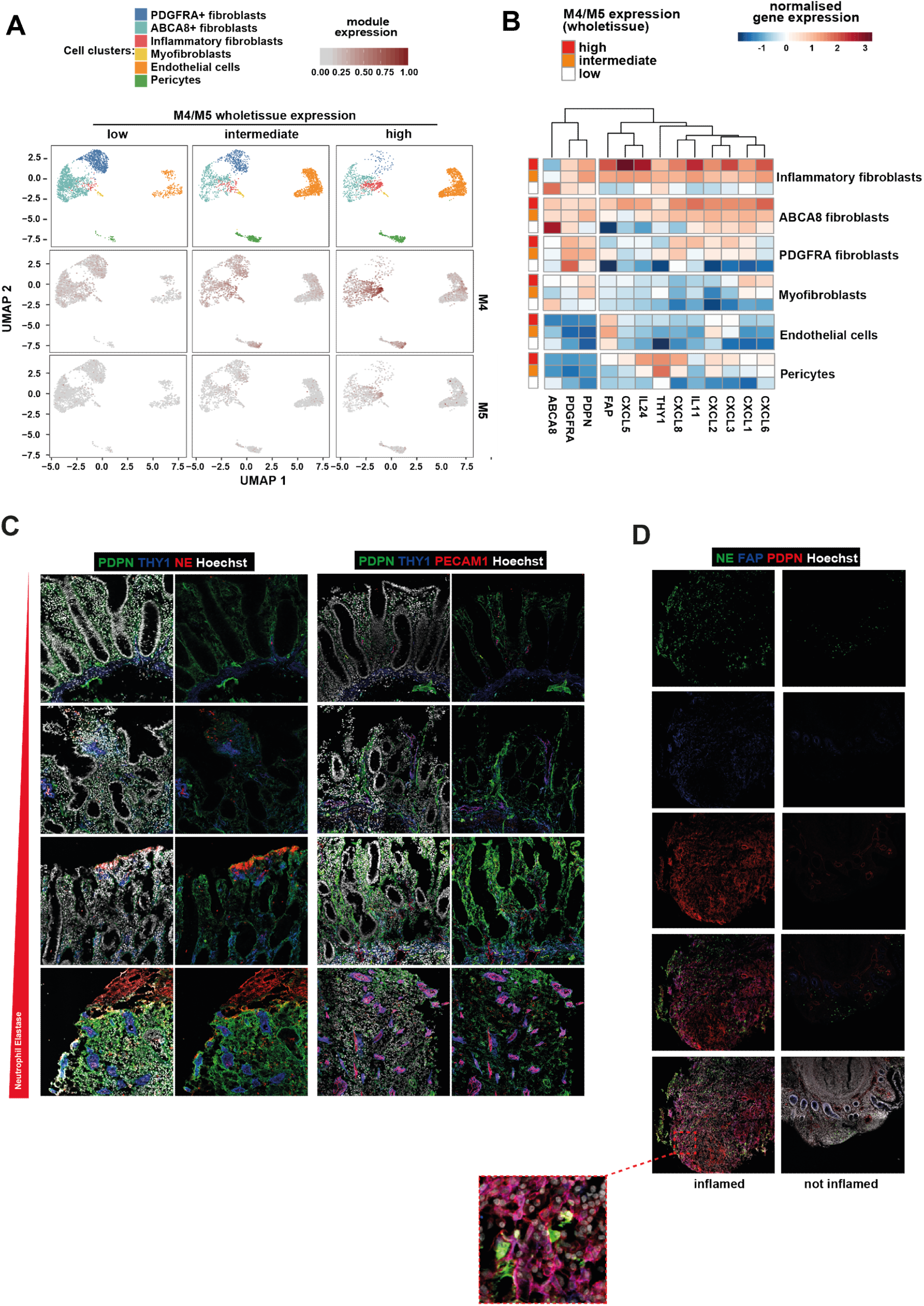
M4/M5 gene expression is associated with neutrophil-attracting fibroblasts and endothelial and perivascular cell expansion. A) UMAP of stromal single-cell profiles showing the different stromal clusters as in Figure 4A for comparison (top panel), and the expression level of M4 (middle panel) / M5 (bottom panel) genes in these clusters (as in Figure 3A). B) Heatmap showing normalised gene expression of the top differentially expressed genes between M4/M5 expression levels within each cell cluster. Expression values are normalised log2 fold changes (Wald statistic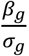)from DESeq2 analyses (see STAR Methods). C) Staining of NE or PECAM1 (red), THY1 (blue) and PDPN (green) in IBD patient tissues with varying grades of neutrophil infiltration. D) Staining of NE (green), FAP (blue) and PDPN (red) in paired inflamed (deep ulcer) and uninflamed IBD tissue.

We confirmed these findings at the protein level, where we found that areas of increased PDPN and THY1 staining were also characterised by a reduced staining of ABCA8 and PDGFRa on fibroblasts (Extended Data Figure 4G). Similarly, we verified that an increased neutrophil presence is associated with more intense PDPN staining and the expansion of the vascular compartment (THY1, CD31), by staining for these markers in different IBD tissues with various grades of neutrophil infiltrates and epithelial damage (Figure 5C). In line with the neutrophil-attracting signature of inflammatory fibroblasts, tissues with dense neutrophil infiltrates (NE+ cells) exhibited the highest level of PDPN on fibroblasts, particularly in areas of profound epithelial cell loss (i.e., ulceration) (Figure 5C). This was associated with the expansion of THY1+ perivascular cells and blood endothelial vessels (PECAM1+THY1+PDPN-), while lymphatic endothelial cells (PDPN+PECAM1+) were mostly absent in areas of neutrophil presence and deep ulceration (Figure 5C). Furthermore, immunofluorescent localisation revealed that in particular PDPN+ fibroblasts that co-expressed FAP (magenta) are located in areas of NE+ neutrophil influx (Figure 5D).

Dissection of the changes within the stromal compartment revealed that the neutrophil infiltrates observed in M4/M5 high patients are associated with the activation of a neutrophil-chemoattractant program in PDPN+FAP+ inflammatory fibroblasts, as well as with angiogenesis and perivascular niche expansion.

### Activated inflammatory fibroblasts drive neutrophil recruitment through IL-1R signalling with high levels of IL-1β at sites of ulceration

To identify potential novel therapeutics targets in the M4/M5 high non-responsive pathotype, we aimed to identify upstream cytokine signalling pathway(s) controlling the observed activation of the neutrophil-attractant program in inflammatory fibroblasts. We culture-expanded primary stromal cell lines (n=33) from surgically resected intestinal tissue of IBD patients and stimulated them with a panel of cytokines associated with IBD ^4^. Of the cytokines assessed, only the NF-kB activators IL-1β and TNF-α, but not IL-6 or OSM, were capable of inducing the expression of CXCL5 in primary stromal cell lines after 3 hours of stimulation (Extended Data Figure 5A). Furthermore, RNA sequencing showed that IL-1β and TNF-α strongly and significantly induced gene expression of all neutrophil-tropic CXCR1 and CXCR2 ligands in fibroblasts, namely *CXCL1, CXCL2, CXCL3, CXCL5, CXCL6* and *CXCL8* (Extended Data Figure 5B). In addition, both cytokines induced the inflammatory fibroblast markers PDPN and FAP (Extended Data Figure 5B). Although TNF-α and IL-1β both induced a chemokine response, the latter was 100-fold more potent (Extended Data Figures 5B + C).

To confirm that the IL-1 signalling pathway is the functionally relevant one for inducing the inflammatory fibroblast phenotype in patients, we developed an *ex vivo* assay using surgically resected tissue from IBD patients. Briefly, we produced conditioned media (CM) from single-cell suspensions of enzymatically digested intestinal tissue. When applied to cultured intestinal fibroblasts, this CM was capable of inducing a robust chemoattractant program (Figure 6A). To determine upstream cytokines driving this response, we blocked IL-1 signalling with the IL-1 receptor (IL-1R) antagonist anakinra (Kineret) or TNF signalling with the anti-TNF agent adalimumab (Humira) in CM. Strikingly, only IL-1R, but not TNF, signalling blockade was able to reduce fibroblast activation in this assay (Figure 6A). These findings demonstrated that soluble mediators contained in gut-resident cell populations of inflamed IBD tissue activate the neutrophil-attracting fibroblast program and that this response is IL-1R but not TNF-dependent. Furthermore, single-cell sequencing showed that inflammatory fibroblasts from M4/M5-high IBD patient tissue were the cell population which demonstrated the strongest IL-1 response pattern (Figure 6B, see Supplementary Table 12 for IL-1 gene expression response), suggesting that this pathway may be associated with the poor therapy response observed in these patients. In line with this, inflammatory fibroblasts and ABCA8 fibroblasts demonstrated the highest fold changes of IL1-receptor (*IL1R1)* expression in M4/M5 high IBD patients (Extended Data Figure 5D). By contrast, TNF receptors (*TNFR1* and *TNFR2*) did not demonstrate this trend (Extended Data Figure 5D). Consistent with the predominant role of IL-1 in these patients, module M5 demonstrated a high enrichment of genes assigned to the inflammasome pathway (Extended Data Figure 5E). Finally, immunohistochemical staining revealed that IL-1β is localised specifically to the ulcer bed and granulation tissue (Figure 6C, top panel), but not uninflamed tissue or tissues where lymphoid aggregates dominate (Figure 6D, top panel). Areas of intense IL-1β labelling also demonstrated intense staining of FAP (Figures 6C and D, bottom panels), suggesting that IL-1R signalling in the ulcer bed is driving the inflammatory fibroblast program characterised by FAP expression.

**Figure 6.**
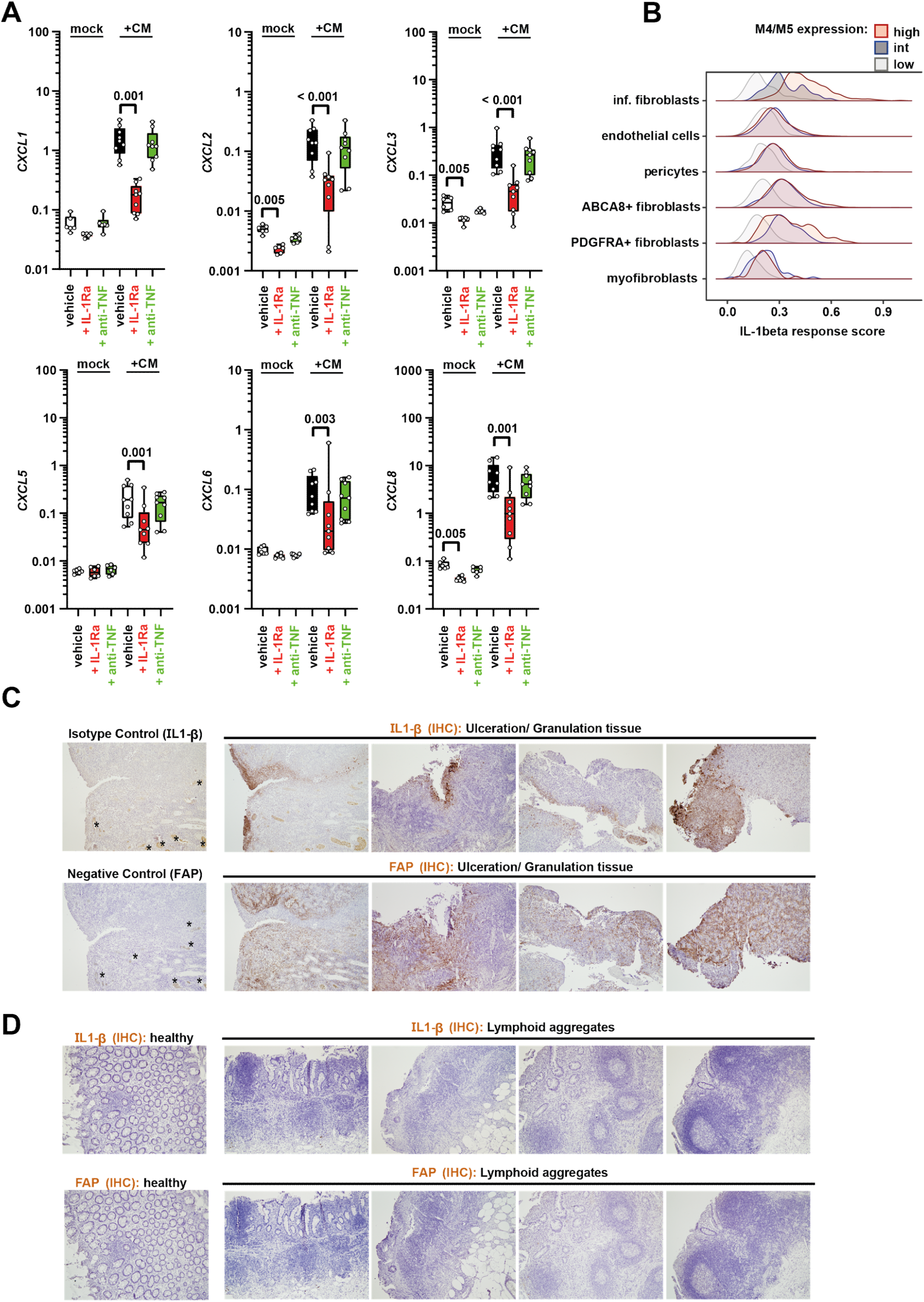
Activated inflammatory fibroblasts drive neutrophil recruitment through IL-1R signalling with high levels of IL-1β at sites of ulceration. A) Ccd18-co fibroblasts were stimulated for 3 h with either mock control or conditioned media produced from IBD patient tissue digests (CM), without pre-treatment (vehicle = PBS), or pre-incubated with IL-1Ra (anakinra) or anti-TNF (adalimumab). Adjusted P-values are shown if significant (p<0.05), Friedman test for paired samples. B) Projection of the IL-1 cytokine stimulation response of Ccd18-co fibroblasts onto stromal cell clusters detected by scRNAseq (see Figure 3A). Score was computed as mean z-score of IL-1 upregulated genes. C) IHC stainings of IL-1β or FAP (DAB, counterstain hematoxylin) in inflamed tissue sections of IBD patients with prominent ulceration and/or granulation tissue. * indicates non-specific staining of erythrocytes or platelets in vessels. D) Stainings as in C), but in inflamed sections of IBD patients with dominant lymphoid aggregates.

Overall, these results identify IL-1R signalling as a key driver of the inflammatory fibroblast/neutrophil recruitment phenotype that is observed in IBD tissues with the high M4/M5 pathotype, which is in turn associated with non-response to multiple therapies currently in use.

## Discussion

Here we integrated transcriptomics, cellular profiling, histopathology and functional assays to identify new, distinct, inflammatory pathotypes associated with therapy non-response in IBD. Non-response to multiple current therapies was associated with a pathotype defined by two gene expression modules that represented IL-1R-dependent inflammatory fibroblast activation, neutrophil accumulation and (peri-) vascular niche expansion at sites of epithelial depletion and deep ulceration. We also identified an additional pathotype associated with an increased presence of lymphoid aggregates that was only linked to patient response to anti-TNF. Combined, these results highlight the existence of distinct pathotypes within the heterogeneous cellular landscape of inflamed tissues in treatment-refractory IBD that are associated with specific treatment outcomes. This provides a novel platform for personalised, precision targeting of existing medications and novel therapeutic targets where current options fail.

Our results highlight neutrophils as a major component of the M4/M5 signature associated with multiple therapy non-response in IBD. In other studies, analysis of tissue-level expression signatures suggested a link between neutrophils and therapy non-response in IBD ^14,15^. Here we have extended those studies by mapping gene expression signatures to intestinal neutrophils isolated from IBD lesions and localising those cells to distinct tissue niches in the inflamed intestine. It is also notable that a dominant neutrophil contribution to the biology of anti-TNF therapy resistance is missing from previous single cell sequencing studies as neutrophils were not analysed ^12 11^. It is not known whether neutrophil accumulation is a cause or a consequence of the chronic tissue damage at sites of tissue ulceration. However, there is evidence that neutrophils can contribute to chronic inflammation through production of extracellular traps (NETs) and the liberation of reactive-oxygen species (ROS) ^27^. We found neutrophils are also the major source of *OSM* expression, a cytokine previously associated functionally with non-response to anti-TNF therapy in IBD ^10^.

The accumulation of neutrophils, activation of fibroblasts and vascular remodelling in response to epithelial damage observed in treatment-refractory IBD lesions is reminiscent of wound healing mechanisms ^28^. It is tempting to speculate that in a subset of non-responsive IBD patients such a chronic wound is a result of an unsuccessful attempt to rebuild the epithelial barrier. Without proper resolution, that process becomes pathogenic, analogous to the concept of a “wound that does not heal” that emerged from the cancer field ^29^. Our single cell RNA sequencing analysis of the stromal compartment identified PDPN and FAP as two markers of fibroblast activation that allowed us to localise inflammatory fibroblasts around ulcers and in proximity to neutrophils. We hypothesise that, rather than being a specialised fibroblast subset, inflammatory fibroblasts may represent an activation state of either ABCA8 fibroblasts residing in the lamina propria of the intestine, or subepithelial PDGFRA fibroblasts. The origin of inflammatory fibroblasts may dependent on the site where damage occurs, i.e. at the epithelial lining layer or deeper into the lamina propria. The very specific localisation of IL-1β in the ulcer bed in proximity to areas of epithelial cell damage suggests disruption of the epithelial cell barrier may be a primary event. Danger-associated molecular patterns (DAMPs) released by necrotic epithelial cells could also lead to the activation of inflammasome pathways and consequently the release of IL-1β. Indeed, several genes associated with an inflammasome signature were found in module M5 of our discovery cohort supporting the idea that inflammasome activation is an upstream event. Early responders to damage at the barrier may also include resident MNP, that can produce excessive IL-1β and IL-23, particularly in the context of IL-10 pathway deficiency ^16^. IL-1R-mediated fibroblast activation leads to the expression of neutrophil-attracting chemokines amongst other inflammatory mediators. Neutrophils are then recruited in high numbers to the site of damage, further contributing to the production of IL-1β in the ulcer bed. The alarmin IL-1α may similarly contribute to the activation of fibroblasts and initiation of colitis ^30 31^, and can be released by necrotic epithelial cells in IBD ^32^. Further studies are required to establish if the IL-1R-driven activation of inflammatory fibroblasts identified here is dominated by IL-1α or IL-1β signalling or both. We did not find IL-18 to be increased in the tissue of M4/M5-high patients.

Currently, sub-categories of IBD are classified by high level phenotypic observations. A lack of knowledge about the molecular pathotypes in IBD means that therapies are currently not prescribed based on the underlying biologic processes they target and therefore often fail. A number of recent studies have tried to address the challenge of therapy non-response by analysing the cellular and molecular network in treatment-refractory IBD. Whilst several genes found in our M4/M5 modules have been previously associated with non-response to anti-TNF therapy or corticosteroids ^10-12,14-17,20^, none of those studies addressed the heterogeneity of molecular inflammatory phenotypes in IBD. By relating molecular signatures (modules) to histologic features, we were able establish this link and identified at least two distinct pathotypes with important implications for patient stratification for therapeutic targeting. In addition to patients with high tissue expression of M4/M5 and substantial ulceration (22/11% of patients in the discovery/early-onset cohorts), we also identified patients with high M6 tissue expression (26% of patients in the discovery cohort) that is associated with increased lymphoid aggregates; the high tissue expression of *CCL19/CCL21/CXCL13* also suggests that this pathotype reflects the presence of fibroblastic reticular-like cells ^25^, as opposed to the inflammatory fibroblast phenotype detected in M4/M5 high tissues. The expression of M6 was very low in the early-onset cohort of paediatric UC and CD. This may reflect the different nature of the samples analysed in the two studies. The latter used endoscopic punch biopsies (mucosa), as opposed to full thickness (mucosa/muscularis/submucosa) samples from surgical specimens in our discovery cohort. Although present in the mucosa, lymphoid aggregates are more prominent in submucosal regions. This requires consideration when interpreting lymphoid tissue signals from endoscopic biopsies of the gut. Tissues high in M6 showed elevated expression *NR3C1* and *ITGA4*, but not *TNF*. This is consistent with our findings that M6 is predictive of non-response to anti-TNF, but not of non-response to corticosteroid or anti-integrin, suggesting that such M6 high patients may benefit from vedolizumab or corticosteroids.

By contrast patients with UC or CD whose tissues show a high M4/M5 signature and ulceration express high amounts of *IL1B* but not *NR3C1, ITGA4 or TNF* suggesting that subgroup may benefit from blocking IL-1R instead of TNF, to target the neutrophil-attractant program in fibroblasts. Indeed, TNF has been shown to promote mucosal healing ^33^ and therefore may be deleterious in patients with deep ulceration that require wound healing. Genetic defects in the IL-1 pathway have been linked to anti-TNF non-response ^34^ and the principle of ameliorating acute intestinal inflammation by blockade of IL-1 signalling has been demonstrated in several pre-clinical models ^35-37^. In case studies of Mendelian disease-like IBD (MD-IBD) with IL-10 deficiency, the blockade of IL-1 signalling has been successfully applied to treat intestinal inflammation ^38,39^, providing proof of concept. Surprisingly, larger scale studies of IL-1 blockade in polygenic IBD patient cohorts are lacking, although trials in acute severe ulcerative colitis are in progress ^40^. Future trials may benefit from stratifying participants for inclusion based on the observations presented here. By dissecting IBD patient heterogeneity at a cellular and molecular level, we provide a rationale for targeting therapeutics to the underlying pathologies, based on histologic features and molecular signatures rather than high-level phenotypic diagnoses.

Our discovery cohort of surgical resection samples from patients with UC or CD highlights the heterogeneity of inflammatory lesions in this difficult-to-treat patient group. This data is just a snap-shot and does not inform on the evolution and dynamics of these distinct pathotypes. However, the presence of M4/M5 signature high patients before treatment in a number of prospective cohorts suggests that deep ulceration and high M4/M5 signature can occur independently of therapy failure. Our study does not address whether lymphoid aggregates and ulceration are independent processes or connected states. Notably, the presence of M4/M5 and M6 is not mutually exclusive, and a small number of tissues exhibited both ulceration and lymphoid aggregates. Further understanding of the natural history of these distinct pathotypes and their relationship to disease dynamics will require longitudinal analyses.

In summary, our combinatorial approach, integrating data across biological levels, identifies new tissular IBD pathotypes that are defined by different molecular, cellular and histopathologic features that are linked to patient responses to current therapeutics. These stratifications provide a basis for personalised targeting of existing medicines and indicate that IL-1 signalling blockade may benefit those individuals with deep ulceration who do not respond to current therapeutics. This may improve treatment trajectories for patients with IBD, both by hastening administration of appropriate interventions and providing a novel mechanism to target in an area of current unmet clinical need.

## Acknowledgements

M.F. has been supported by the Roche postdoctoral fellowship program (2016-2017) and is currently supported by the Oxford-UCB-prize fellowship scheme. M.P. has been supported by the Roche postdoctoral fellowship program (2017-2019). M.A.J. is funded by the Kennedy Trust for Rheumatology Research and supported by a Junior Research Fellowship from Linacre College Oxford. The authors thank members of the Oxford TGU biobank, especially James Chivenga, Adam Isherwood and Roxanne Williams, as well Gkentiana Zavalani for facilitating access to patient samples. This work was supported by the National Institute for Health Research (NIHR) Oxford Biomedical Research Centre (BRC). The views expressed are those of the author(s) and not necessarily those of the NHS, the NIHR, or the Department of Health. Parts of this work were supported by the BRC3 mucosal immunology response mode funding and the Wellcome Trust (212240/Z/18/Z). The authors further thank Johanna Pott for helping to set up *ex vivo* functional assays and the following Kennedy Institute core facility members for excellent technical support and training: Jonathan Webber, Bryony Stott, Ida Parisi, Claudia dos Santos Duarte and Volodymyr Nechyporuk-Zloy.

## Author contributions

Conceptualisation, M.F., M.P., M.A.J and F.M.P. Methodology, M.F., M.P., M.A.J., I.K., S.B., K.R., D.S., M.A., K.W., G.W., Roche Fibroblast Network Consortium, A.E., S.S., S.R., S.P.T. and F.M.P. Software, M.A.J. and I.K. Validation, M.F., M.P., M.A.J. and I.K. Formal Analysis, M.F., M.P., M.A.J., I.K., K.R., E.C. and A.E. Investigation, M.F., M.P., S.B., D.S., H.S, R.R. and A.G. Resources, M.F., M.P., Z.C., R.R., R.S.P., E.H.M., A.G., T.T., Oxford IBD Cohort Investigators, Roche Fibroblast Network Consortium, S.M., L.T., S.S., S.R., S.P.T. and F.M.P. Data Curation, M.F., M.P., M.A.J., I.K., S.B., K.R., Z.C., D.S., R.R., L.T. and A.E. Writing –Original Draft: M.F., M.P., M.A.J., and F.M.P. Writing – Review & Editing: all authors listed. Visualisation, M.F., M.P., M.A.J., I.K., and K.R. Supervision, M.F., M.P., S.B., S.S., S.R.,S.P.T. and F.M.P. Project Administration, M.F., M.P., TGU Biobank Consortium, S.P.T. and F.M.P. Funding Acquisition, M.F., I.K., S.B., Z.C., S.R., S.P.T. and F.M.P.

## Competing Interests Statement

*Fiona Powrie, PhD, FRS, FMedSci*

Grants/Research support: Roche and Janssen

Consulting fees: GSK and Genentech

*Simon Travis, DPhil FRCP*

Grants/Research Support: AbbVie, Buhlmann, Celgene, IOIBD, Janssen, Lilly, Pfizer, Takeda, UCB, Vifor and Norman Collisson Foundation.

Consulting Fees: AbbVie, Allergan, ?biomics, Amgen, Arena, Asahi, Astellas, Biocare, Biogen, Boehringer Ingelheim, Bristol-Myers Squibb, Buhlmann, Celgene, Chemocentryx, Cosmo, Enterome, Ferring, Giuliani SpA, GSK, Genentech, Immunocore, Immunometabolism, Indigo, Janssen, Lexicon, Lilly, Merck, MSD, Neovacs, Novartis, NovoNordisk, NPS Pharmaceuticals, Pfizer, Proximagen, Receptos, Roche, Sensyne, Shire, Sigmoid Pharma, SynDermix, Takeda, Theravance, Tillotts, Topivert, UCB, VHsquared, Vifor, Zeria.

Speaker fees: AbbVie, Amgen, Biogen, Ferring, Janssen, Pfizer, Shire, Takeda, UCB. No stocks or share options.

*K*.*G. L. and A*.*P*.*F*. are employees of Roche Ltd. All authors declare no commercial or financial conflict of interest.

## Online Methods

### Patient cohorts and ethics

Patients eligible for inclusion in the discovery cohort were identified by screening surgical programs at Oxford University Hospitals. Samples were obtained from patients undergoing surgical resection of affected tissue for ulcerative colitis (UC), Crohn’s disease (CD) or colorectal cancer (CRC) (used as non-IBD controls). All tissue samples included in the study were classified by pathological examination as either macroscopically active inflamed or uninflamed. Additional samples were also obtained from CD and UC patients or from healthy individuals by biopsy. All patients gave informed consent and collection was approved by NHS National Research Ethics Service under the research ethics committee references IBD 09/H1204/30 and 11/YH/0020 for IBD or GI 16/YH/0247 for CRC samples and gut biopsies from healthy individuals. Samples were immediately placed on ice (RPMI1640 medium) and processed within 3 hours. All patients gave informed consent and data was fully anonymised prior to analyses. For replication of prospective findings in the discovery cohort, public datasets were used that were derived from endoscopic tissue samples of IBD patients ^15,20,21,41^ (GSE16879, GSE73661, GSE109142, GSE57945).

### Isolation of cells from tissue and blood samples

After removing external muscle and adipose layers, and removing bulk epithelial cells by repeated washes in PBS containing antibiotics (Penicillin-streptomycin, amphotericin B, gentamicin, ciprofloxacin) and 5mM Ethylendiaminetetraacetic acid (EDTA, Sigma Aldrich), tissue from surgical resections was minced using surgical scissors. In the case of endoscopic biopsies, the epithelial wash was omitted. Minced tissue was subjected to multiple rounds of digestion in RPMI1640 medium containing 5% fetal bovine serum (FBS), 5mM HEPES, antibiotics as above, and 1mg/ml Collagenase A and DNase I (all from Sigma Aldrich). After 30 minutes, digestion supernatant containing cells was taken off, filtered through a cell strainer, spun down and resuspended in 10ml of PBS containing 5% BSA and 5mM EDTA. Remaining tissue was then topped up with fresh digestion medium until no more cells were liberated from the tissue.

### Primary culture expansion and conditioned media production

Primary stromal cell lines were expanded by plating the single-cell suspension of tissue digests onto plastic cell culture vessels and expanding the adherent fraction (>95% CD45-EPCAM-CD31-cells, not shown) in RPMI1640 (with 20%FCS, antibiotics, 5mM HEPES) (Sigma). Primary cell lines were used for assays between passage number 7 and 15. For the production of conditioned media, sorted cell populations were plated at 1.000.000 cells/ml in cell culture dishes and RPMI1640 containing 5%FCS (LifeTechnologies), antibiotics, and 5mM HEPES for 16 hours. After that, supernatants were aspirated, spun down to remove cells, and frozen at -80°C until further use.

### Fluorescence-activated cell sorting (FACS) and analysis

Single-cell suspensions obtained from tissue digests were stained for FACS analysis or sorting with antibodies (all from Biolegend, except anti-Pdpn: clone NZ-1.3 from eBioscience) in PBS with 5% BSA and 5mM EDTA for 20 minutes on ice. After washing in the same buffer, cells were analysed (LSRII or Fortessa X20) or sorted (Aria III, 100um nozzle).

### *Ex vivo* assay of IBD patient conditioned media transferred onto fibroblasts

For the stimulation of stromal cells, either Ccd18-Co colonic fibroblasts (ATCC #CRL-1459) or primary stromal cell lines (isolated as above) were plated at 20000 cells/well in a 48-well plate. Plated cells were starved for 72 hours in culture medium without FCS, before stimulation with cytokines or conditioned media (pre-diluted 1:3 in starving medium) for 3 hours at 37°C. For blockade experiments, recombinant cytokines in starving medium or conditioned media were pre-incubated with 2mg/ml Anakinra (Kineret) or Adalimumab (Humira) for 1 hour at RT (shaking) before stimulation of cells. After 3 hours, supernatants were taken off and cells lysed directly in appropriate RNA lysis buffer.

### Isolation of RNA from tissue samples and cell populations

Endoscopic punch biopsies or dissected tissue pieces from surgical resections were stored in RNAlater (Qiagen) upon collection until further processing. Tissue was homogenised using the soft tissue homogenizing CK14 kit (Precellys, Stretton Scientific 03961) in 300µl of RLT lysis buffer (Qiagen) and 20µM DTT (Sigma). RNA was isolated using the Qiagen Mini kit with a DNA digestion step (Qiagen). Bulk-sorted cell populations and cultured cells were directly lysed in RNA lysis buffer, followed by RNA isolation with the according kits and on-column DNase treatment.

### Sequencing of RNA from whole tissue and sorted cell populations

Sequencing libraries were prepared using either the QuantSeq 3’ mRNA-Seq FWD Library Prep Kit (Lexogen) for whole tissue samples or the Smart-seq2 protocol ^42^ for bulk and cultured cell populations (with our own in-house indexing primers). Libraries were sequenced using an Illumina HiSeq4000 with 75bp paired-end sequencing ^43^. For qPCR analysis, 15 to 250ng of RNA was reverse-transcribed using the High-Capacity cDNA Reverse Transcription Kit (Applied Biosystems) and qPCR performed using Precision Fast qPCR mastermix with ROX at a lower level, 12.8mL (Primer design, Precision FAST-LR), and Taqman probes (Life Technologies).

Bulk RNA sequencing data were analysed using the bulk processing aspect of pipeline_scrnaseq.py (https://github.com/sansomlab/scseq). Data quality was assessed using pipeline_readqc.py (https://github.com/cgat-developers/cgat-flow). Sequenced reads were aligned to the human genome GRCh38 using Hisat2 (version 2.1.0) ^44^ using a reference index built from the GRCm38 release of the mouse genome and known splice sites extracted from Ensembl version 91 annotations (using the hisat2_extract_splice_sites.py tool). A two-pass mapping strategy was used to discover novel splice sites (with the additional parameters: -- dta and --score-min L,0.0,-0.2). Mapped reads were counted using featureCounts (Subread version 1.6.3; Ensembl version 91 annotations; with default parameters) ^45^. Salmon v0.9.1 was used to calculate TPM values ^46^ using a quasi-index (built with Ensembl version 91 annotations and k=31) and gc bias correction (parameter “--gcBias”). For heatmap visualisations of gene expression levels, z-scores of TPM values and Manhattan distances were calculated within the *heatmap2* package in R. Differential expression analyses were performed using DESeq2 (v1.26.0)^47^. Raw data will be deposited in GEO.

Pathway enrichment analysis for groups of genes associated with cell types was carried out by the *enrichGO* function from the *clusterProfiler* package in R ^48^. “Cellular component” GO annotation terms were used as pathways.

### Identification and quantification of gene co-expression modules in discovery data

To reduce dimensionality within the dataset, an unbiased approach was used to collapse genes with similar expression patterns in the discovery RNAseq dataset. Normalised (TPM) counts were considered for all genes across all samples, including both inflamed and uninflamed tissues from the IBD patients and the samples from the CRC controls. These were filtered to remove genes with zero counts in over half of the samples and log transformed following the addition of a pseudo count. Transformed counts were then used to define modules of correlated genes using the weighted gene co-expression network analysis approach (WGCNA) in R ^49^. In brief, this process calculates pair-wise Pearson correlation estimates between all genes. These are then raised to the power of a soft-threshold, in this case raising correlation coefficients to the power of 9, which magnifies the differences between large and small correlations. Finally, the network of these amplified correlations (where each gene is a node and each edge is a correlation) is used to generate a topological overlap matrix (TOM). This represents the similarity of expression patterns between a given pair of genes in the data set, similar to the correlation matrix, but taking into account their shared correlation with other genes. Finally, hierarchical clustering of the TOM is used to assign genes into modules based on their co-expression pattern. The *pickSoftThreshold* function was used to identify 9 as an appropriate soft-threshold. The *blockwiseModules* function was then used with this threshold to automatically carry out the aforementioned process and assign genes to modules. Parameters for the function were as follows, a minimum module size of 30 genes, a mergeCutHeight of 0.1, reassignThreshold of 0, and using a signed network.

The resultant module definitions were quantified using the eigengene approach within WGCNA. An eigengene is a quantitative representation of the expression of a module as a whole and is derived from the first component of a principle components analysis restricted to the expression data of just the genes in the module. Eigengenes for the modules defined in the resection data were calculated using the *moduleEigengenes* function.

Correlations between clinical and metadata measures and module eigengenes were assessed using Pearson correlations with p-values estimated using the *corPvalueStudent* function and adjusted for multiple testing. Benjamini-Hochberg correction using the *p*.*adjust* function was used for all analyses with adjusted p-values. This was carried out on the inflamed IBD tissue samples and CRC tissue samples combined and also on the inflamed IBD tissue samples alone. Eigengenes were also compared between paired inflamed and uninflamed tissues sections using a t-test, adjusting for multiple testing across modules.

Cell type composition scores were estimated for each resection sample using the *xCellAnalysis* function from the xCell package ^19^. Correlations between module eigengenes and the derived cell type scores were visualized for all cell types scored in over 25% of samples and used to infer the cell types represented by modules within the whole tissue data (discovery cohort).

### Quantifying module associations with clinical variables in replication datasets

Publicly available RNAseq (^15,41^ (GSE57945, GSE109142)) or microarray data ^20,21^ (GSE16879, GSE12251) were downloaded from the NCBI gene expression omnibus. These were pre-existing enumerated gene counts in the case of the RNAseq datasets and raw array data in the case of the microarray sets. The latter were processed and normalised to gene counts using the *rma* function from the affy package ^50^, summing values for probes associated with the same gene symbol. Across all datasets, gene symbol annotations were used to map the genes to the module assignments generated from the discovery resection tissue dataset, dropping genes that were not observed in the replication dataset under consideration. The percentage of genes missing from the original module definitions was recorded but was generally low across all datasets. Mapped module assignments were then used to generate eigengenes from the replication expression datasets using the *moduleEigengenes* function. Correlations between clinical metadata and eigengenes in replication datasets was performed using Pearson correlations as for the discovery dataset. In the case of the paediatric cohort data (GSE57945), Mann-Whitney U tests were used to compare the modules between patients scored as ulcerated or not in metadata and hierarchical clustering used to group patients based on M4,5, and M6 expression as for the discovery cohort.

Differences in pre-treatment module eigengene values between responders and non-responders in prospective studies were assessed using Mann–Whitney *U* tests, adjusting p-values for testing of multiple modules within each dataset. In the Haberman et al. 2019 study we only considered patients on corticosteroid therapy, combining both patients that received oral and intravenous administration. In the case of the Arjis 2018 study, which tested multiple different therapies and treatment regimens, we used ANOVA to identify any differences between responders and non-responders across all combinations adjusting for regimen and used post-hoc Mann-Whitney U tests to identify individual treatment regimens where modules were significantly different by response.

Meta-analysis of the expression of the M4 and M5 modules across responders and non-responders in the various replication datasets was carried out using the meta package in R ^51^. The anti-TNF response data was used from the Arijs 2008 and 2018 papers and the corticosteroid response data was from the Haberman et al. study. Only the 4 week treatment condition was included from the anti-intergrin data from Arijs 2018, as this was the only one that proved significantly different for either M4 or M5. A random effects meta-analysis was carried out comparing standardised mean differences between patient groups using the exact Hedges estimate.

In the prospective cohorts, the predictive value of the expression of single genes for response to treatment was assessed using a simple logistic regression where response was the outcome and gene expression the sole predictor. Modelling was carried out for all genes also observed in the discovery cohort, for each of the prospective studies, using the *glm* function in R. The predictive ability of each gene in each dataset was summarised as the area under the curve (AUC) of a receiver operating characteristic (ROC) curve. AUC values for each gene were generated by applying the *roc* function from the pROC package to predictions generated from the logistic regression models. The relative predictive power of genes within modules of interest was compared by summing the rank of genes (based on their AUC value) across datasets and comparing these cumulative ranks between modules.

### Pathological scoring of histology using the Nancy index

Formalin-fixed paraffin-embedded (FFPE) and hematoxylin & eosin (H&E)-stained tissue sections of IBD patients were scored according to the Nancy index, based on criteria reported in ^18^.

### Immunohistochemistry and quantitative histopathology

Tissue specimens were either fixed for 48 hours in 4% neutral-buffered formalin (Sigma) and embedded in paraffin (“FFPE”), or fixed for 24 hours in 2% PFA in phosphate buffer containing L-lysine and Sodium periodate and frozen in OCT (Sigma) after soaking in 30% sucrose for 48 hours (“OCT”). Freshly cut, dewaxed, and rehydrated FFPE sections (5µm) were subjected to heat-induced antigen retrieval by boiling in Target Retrieval Solution (Dako, pH=6, for all stainings except neutrophil elastase) for 15minutes (microwave). This was followed by 15 minutes of blocking in Bloxall solution (Vector Labs), 60minutes blocking in 5%BSA/TBST with 5% serum of the secondary antibody species (Sigma), and 15minutes of blocking in avidin followed by biotin solution (Vector Labs). All steps were performed at ambient temperature. Tissue sections were incubated with primary antibodies in 5%BSA/TBST overnight (>16 hours) at 4°C. Following incubation, biotinylated or HRP/AP-conjugated secondary antibodies were applied for 2 hours (RT) in 5% BSA/TBST. For biotinylated secondary antibodies, AB complex (Vector Labs) was incubated for another hour in TBST (RT). Chromogenic stains were developed by applying DAB HRP substrate solution (Vector Labs) and counterstained for 5minutes in Hematoxylin solution (Sigma). Slides were then dehydrated and mounted in DPX (Sigma) mounting medium.

Whole section imaging of chromogenic sections was performed on a NanoZoomer S210 digital slide scanner (Hamamatsu). Slide scans of all stains can be made available upon request. Scanned tissue sections, stained using DAB immunohistochemistry, were analysed using Indica Labs HALO® image analysis platform. A consultant gastrointestinal pathologist manually annotated each slide, dividing the mucosa into normal and inflamed. The tissue was scored using Indica Labs analysis modules CytoNuclear v2.0.5, detecting DAB positive and negative cells in inflamed areas. Pathologic features (ulceration/granulation tissue, granulomas, crypt abscess/cryptitis, lymphoid aggregates and architectural distortion/mucin depletion) were manually annotated by a consultant pathologist with a special interest in gastrointestinal pathology. The area of each annotated feature was automatically calculated by the HALO software. Nuclei (cells) in areas of interested and the whole tissue section were detected and counted using Indica Labs – CytoNuclear v2.0.9 analysis module. Scores (%) were normalised to the number of nuclei that were found within a pathological feature over the total number of nuclei detected in the whole tissue section. These normalised counts were used to investigate Pearson correlations between features and correlations with module eigengenes.

10µM thick OCT sections were incubated in blocking buffer (PBS1X, 5% Goat serum, 2% FCS and human FcBlock (Miltenyi) with primary antibodies overnight at 4°C. The next day AF488 Donkey anti Rat, AF647 Donkey anti Goat, AF555 Donkey anti Rabbit or strepatividin-AF568 were applied for 1h at RT in blocking buffer. Finally, nuclei were stained with Hoechst 28332 (Life Technologies) for 15min at RT in blocking buffer and then mounted in ProlongGold mounting medium (Life Technologies) prior to imaging with the spectral detector of a Zeiss confocal LSM 880 microscope.

### Preparation of cells for single-cell RNA sequencing

Four pairs of biopsies were pooled, minced and frozen in 1mL of CryoStor® CS10 (StemCell Technologies) at -80°C then transferred in LN_2_ within 24 hours. Single-cell suspensions from these endoscopic biopsies were then prepared by thawing, washing and subsequent mincing of the tissue using surgical scissors. Minced tissue was then subjected to rounds of digestion in RPM-1640 medium (Sigma) containing 5% FBS (Life Technologies), 5mM HEPES (Sigma), antibiotics as above, and Liberase TL with DNAse I (Sigma). After 30 minutes, digestion supernatant was taken off, filtered through a cell strainer, spun down, and resuspended in 10ml of PBS containing 5% BSA and 5mM EDTA. Remaining tissue was then topped up with fresh digestion medium until no more cells were liberated from the tissue. Cells were then stained and FACS-sorted, as described above for live EPCAM-CD45-cells, before being taken for microfluidic partitioning (see below).

### 10x library preparation, sequencing, and data analysis

Single-cell RNAseq data was generated from disaggregated intestinal tissue sorted for Sorted CD45-EPCAM-stromal cells. Viable cells were subjected to a standard droplet single-cell cDNA library preparation protocol. The experimental details to generate cDNA libraries are described in a separate manuscript (https://www.biorxiv.org/content/10.1101/2021.01.11.426253v1). We demultiplexed FASTQ files for each 10X library using the Cell Ranger (v3.1.0) mkfastq function ^52^. We then mapped reads to the GRCh38 human genome reference using Kallisto ^53^ (v0.46.0) and quantified gene by cell-barcode UMI matrices with Bustools (https://github.com/BUStools/bustools) (v0.39.0).For quantification, we used gene annotations provided by Gencode ^54^ (release 33), keeping only protein coding genes and collapsing Ensembl transcripts to unique HGNC approved gene symbols.

We filtered for potentially empty droplets and damaged cells by excluding droplets with fewer than 500 unique genes and libraries with greater than 20% of reads assigned to mitochondrial genes. We pooled the resulting high-quality cells from each 10X library into a single cell by gene UMI matrix. We normalized for read depth with the standard logCP10K normalization procedure for gene *g* and cell *si*:

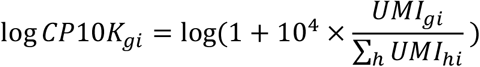

We performed PCA analysis on the top 2000 most variable genes, identified with the VST method implemented in the Seurat ^55^ R package. For PCA, we z-scored each variable gene and computed the top 30 eigenvectors and singular values with the truncated SVD procedure, implemented in the RSpectra (https://github.com/yixuan/RSpectra) R package. We defined PCA cell embeddings by scaling eigenvectors by their respective singular values. To account for potential batch effects in the PCA embeddings, we modelled and removed the effect of 10X library as identified using the Harmony algorithm. For Harmony ^24^, we set the cluster diversity penalty parameter θ to 0.5 and used default values for all other parameters. We evaluated the effect of library mixing before and after Harmony using the Local Inverse Simpson’s Index (LISI), described in the Harmony manuscript ^24^. We evaluated the significance of the LISI change with a t-test, with degrees of freedom equal to the number of libraries minus 1. To visualize the cells in 2 dimensions, we input the Harmonized PCs into the UMAP (arXiv:1802:03426 [stat.ML]) algorithm.

### Identification of marker genes within single-cell RNA sequencing

We performed joint clustering analysis on all scRNAseq libraries using the cells’ Harmonized PCA embeddings. With the 30-nearest neighbour graph, we computed the unweighted shared nearest neighbour (SNN) graph, and truncated SNN similarity values below 1/15 to zero. We then performed Louvain clustering, based on the R/C++ implementation from Seurat, at resolution=0.3, resulting in 8 clusters. We identified upregulated marker genes in each cluster using pseudobulk differential expression with negative binomial regression, implemented in the DESeq2 R package. For pseudobulk analysis, we collapsed cells from the same donor and cluster into one pseudobulk sample, summing the UMI counts from each cell. We then performed differential expression analysis on these pseudobulk samples, with the design *y* ∼ 1 + *cu*. This design assigns each gene an intercept term (i.e. mean expression), a multiplicative offset for each cluster. We addressed the degeneracy of the design matrix by assigning a Gaussian prior distribution to the cluster effects (DESeq2 parameter βPrior=TRUE). The full results for this differential expression analysis are reported in Supplementary Table 9.

### Differential expression analysis of single-cell data by inflammatory status

We performed differential expression to associate genes with inflammation status within each single-cell cluster. We used DESeq2 on the pseudobulk samples described above, this time analysing each cluster separately with the design *y* ∼ 1 + *infsamstsus*. We treated InflamStatus as a random effect (DESeq2 parameter βPrior=TRUE) and recovered a mean multiplicative offset for each of the three inflammatory status categories.

### Single-cell gene set enrichment scoring

Single-cell gene-set enrichment scores were computed for WGCNA modules and cytokine stimulation signatures using the same strategy. For each gene in the gene set, we computed Z scores (mean centred and unit variance scaled) of logCP10K normalized expression across all cells. Then we summed the Z-scores of genes in the gene set to compute a single gene set score for each cell. This procedure is summarized in the formula below, used to compute the score *S*_*G,i*_ for geneset *G* and cell *i* using normalized expression *y*_*gi*_, gene mean *μ*_*g*_, and gene standard deviation *σ*_*g*_.

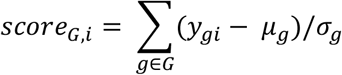

### Single-cell trajectory analysis

We performed trajectory using the principal curve method, implemented in the princurve R package (https://www.jstor.org/stable/2289936). We fit a principal curve to all fibroblasts by inputting harmonized UMAP coordinates into the principal_curve function. This mapped fibroblasts to a non-linear, one dimensional space and assigned each cell a unique position, from 0 to 100, along this trajectory. To directly visualize the abundance of each cluster along the trajectory, we plotted the relative density of each cluster along the trajectory. In these density plots, ABCA8^+^ fibroblasts grouped towards the beginning (position=32) of the trajectory, PDPN^+^ fibroblasts in the middle (position=59), and PDGFRA^+^ fibroblasts towards the end (position=82). This distribution along the trajectory is also reflected by the canonical markers of these populations. To visualize this, we discretized pseudotime by binning into 100 uniform-density windows, chosen so that each window has the same number of cells. We then plotted the scaled gene expression values of ABCA8, PDPN, and PDGFRA, summarized by mean expression (point) and 95% confidence interval (line).

## Data and Code availability

RNA sequencing data will be made accessible via GEO (bulk) and ImmPort (single-cell), and all analysis code will be made available through a GitHub repository at https://github.com/microbialman/IBDTherapyResponsePaper.

**Extended Data Figure 1.**
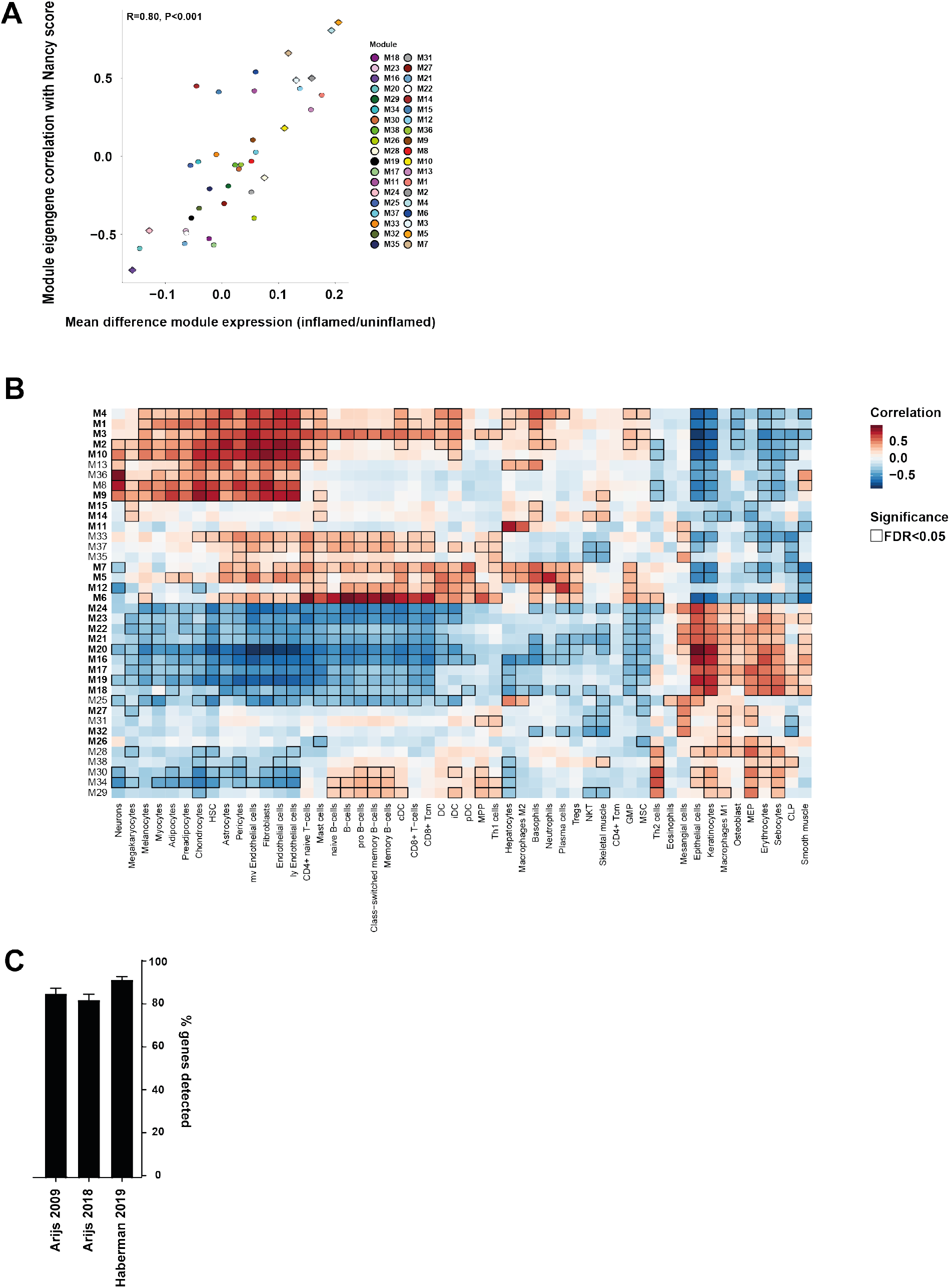
A) Scatterplot of the module expression difference between inflamed and uninflamed tissues paired from the same patients versus the correlation of the module with the Nancy score across all IBD and non-IBD tissues. Points highlighted with a diamond indicate a significant difference in paired t-test between inflamed/uninflamed tissue (FDR p<0.1). B) Heatmap of module eigengene – cell type correlations; cell types were deconvoluted from whole tissue expression data using *xCell*. Modules highlighted in bold were fund to be associated with histologic inflammation. C) Percentage of genes within each module that were detectable in the publicly available datasets. Bars show mean and standard error across all modules for each dataset. ^15,20,21^.

**Extended Data Figure 2.**
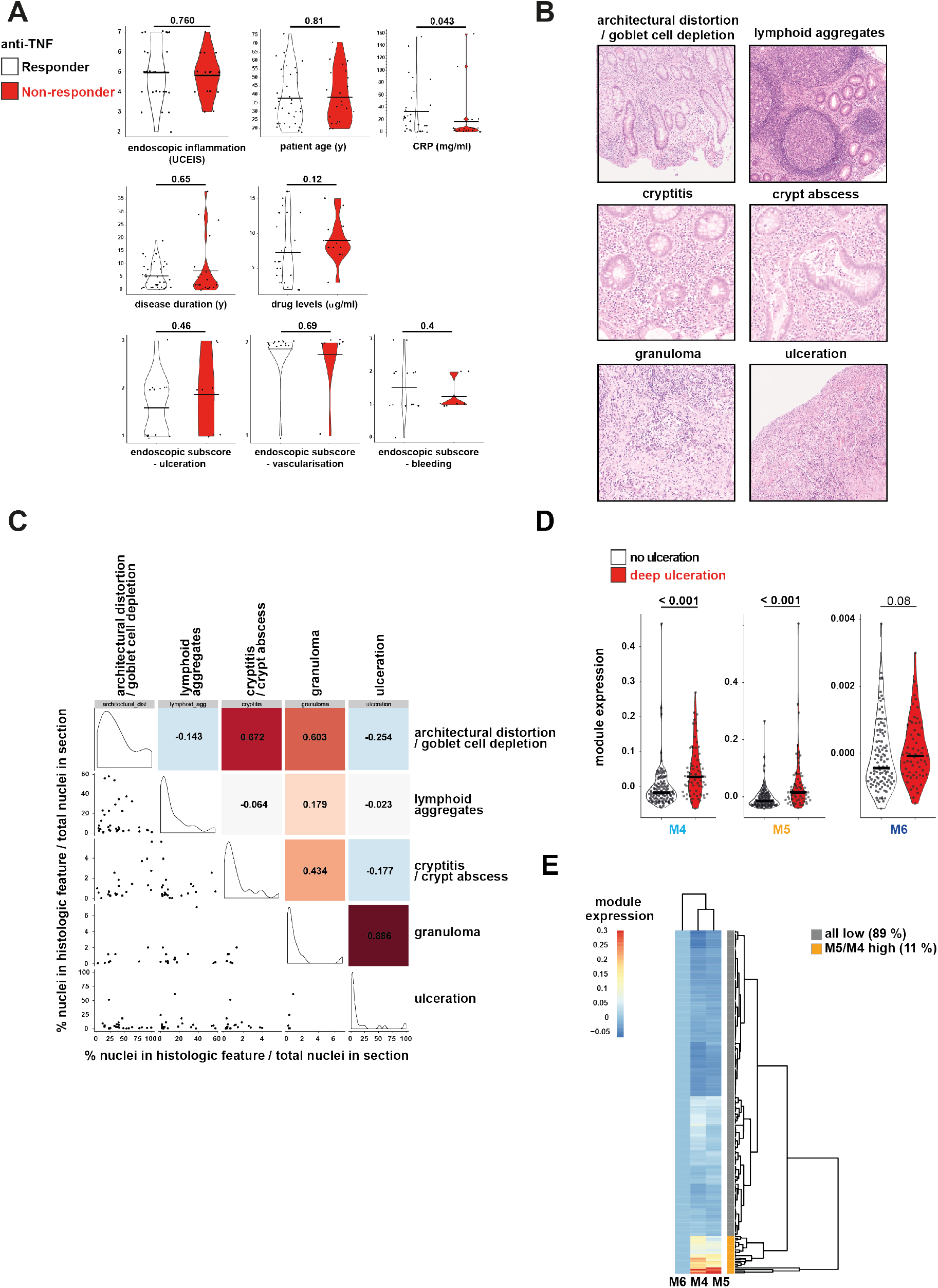
A) Clinical and endoscopic measures in responders and non-responders to anti-TNF therapy before the start of treatment (horizontal bars indicate geometric mean, Wilcoxon signed rank test P values are given). B) Representative images of the various pathological features quantified on H&E histology of resected tissue from IBD patients. C) Correlation plot of histological features, quantified as the % of nuclei within the feature area relative to the nuclei with the total section area. Numbers and colours in upper right corner indicate the Pearson correlation coefficient; histograms on diagonal show the value distribution of the features within IBD patient tissues; scatter plots in the lower left corner show the individual datapoints. D) Violin plots of eigengene expression of M4, M5 and M6 in inflamed tissues of IBD patients with or without deep ulceration observed in a replication cohort of paediatric CD and UC (n=172). E) Classification of M4/M5 high and M4/M5 low patients in the paediatric replication cohort, based on hierarchical clustering on module eigengene values.

**Extended Data Figure 3.**
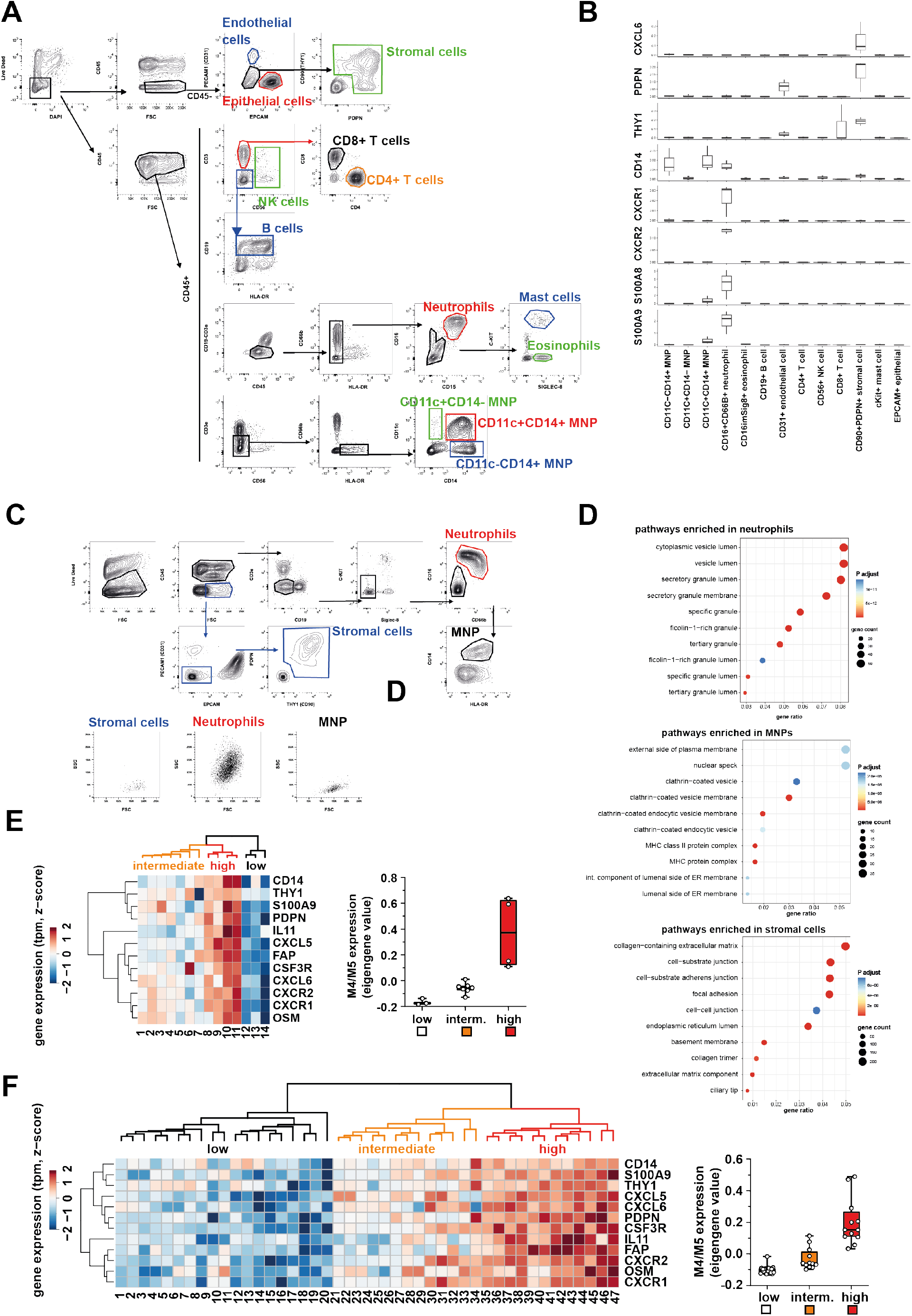
A) Gating strategy for FACS sorting of hematopoietic and non-hematopoietic cell populations from non-IBD and IBD patient tissue. B) Normalised gene expression (qPCR, relative to *RPLP0* expression) of selected genes from M4 and M5 in cell populations sorted as in A. C) FACS-gating strategy for sorting of neutrophils, stromal cells and MNPs from tissue samples of IBD patients. D) Gene set enrichment analysis using Gene Ontology (GO) Cellular Components pathway terms, based on all genes significantly enriched (p adjusted <0.05, |log2 fold change| > 2) in either neutrophils MNPs or stromal cells (Supplementary Table 8). E+F) Heatmaps of whole tissue gene expression of selected genes that are representative (=highly correlative) of M4 and M5 expression (qPCR, z-score transformed gene expression values); unsupervised clustering (Manhattan) distinguishes subgrouping into M4/M5 low, intermediate and high samples; the box-plots on the right show the eigenvalues of all detected genes on a per patient basis. The respective heatmaps refer to tissue samples used for FACS analysis (E) and IHC analysis (F) as shown in Figures 3D, F and G.

**Extended Data Figure 4.**
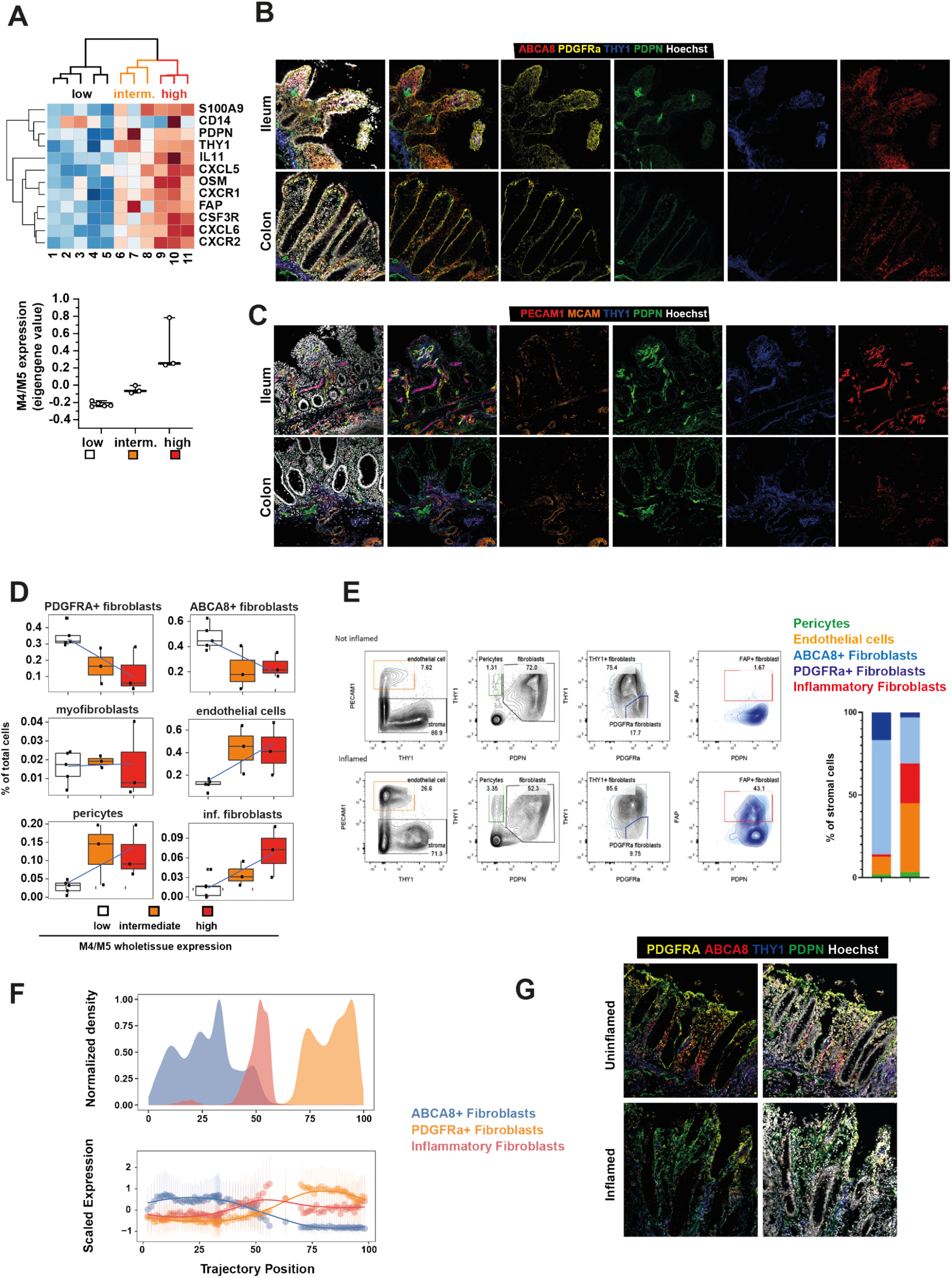
A) Heatmap of whole tissue gene expression of selected genes that are representative of (highly correlated with) M4 and M5 expression (qPCR, z-score transformed gene expression values); unsupervised clustering (Manhattan) groups samples into M4/M5 low, intermediate and high from the set of IBD patients whose samples were profiled by single cell RNA sequencing; the box-plots on the right show the eigenvalues of all detected genes on a per patient basis. B) Immunofluorescent staining of ABCA8 (red), PDGFRA (yellow), THY1 (blue), Podoplanin (PDPN, green) and nuclei (Hoechst, grey) in ileum and colon of resected tissue from IBD patients (not inflamed). C) Immunostaining of PECAM1 (red), MCAM (orange) THY1 (blue), Podoplanin (PDPN, green) and nuclei (Hoechst, grey) in ileum and colonic resected tissue from IBD patients (not inflamed). D) Box plot showing the proportion of the cell types in M4/M5 low, intermediate and high groups, as detected by scRNAseq. E) FACS analysis of live stromal cells (CD45-, EPCAM-) in resected tissue from an IBD patient (adjacent not inflamed and inflamed tissue). Gates for endothelial cells (PECAM1+), Pericytes (THY+, PDPN-), ABCA8+fibroblasts (THY1 high, PDGFRa low), PDGFRA+ fibroblasts (PDGFRA high, THY1 low) and inflammatory fibroblasts (FAP+) are shown. F) Pseudotime analysis of ABCA8+, PDGFRA+ and inflammatory fibroblasts in the single-cell dataset. Cell densities (top row) or canonical markers (bottom) are shown along the trajectory, binned to 100 uniform-density windows (each window has the same number of cells). G) Representative immunofluorescent stainings of PDGFRA (yellow) and ABCA (red) staining on fibroblasts in paired inflamed and uninflamed samples of the same IBD patient.

**Extended Data Figure 5.**
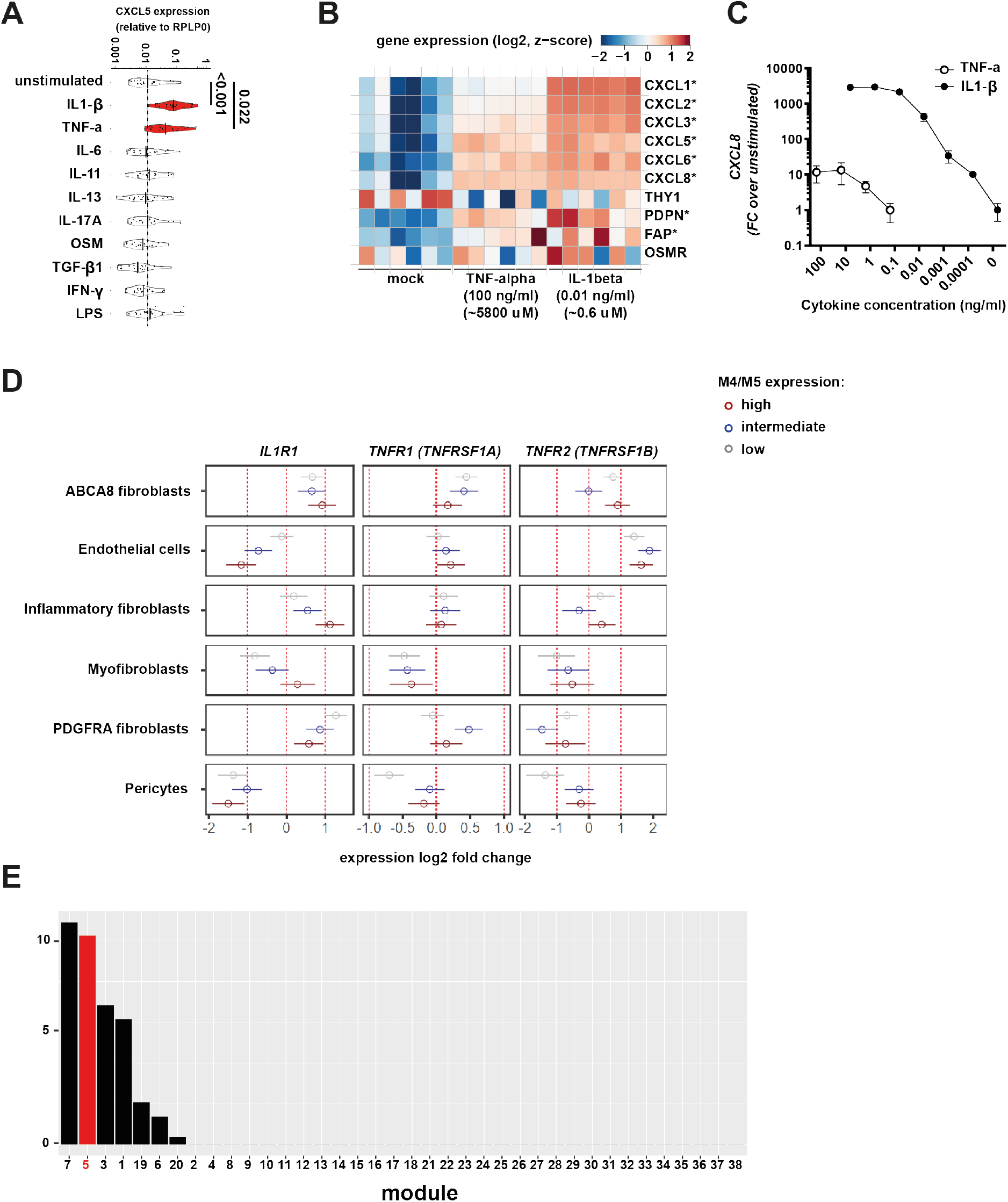
A) Primary fibroblast cell lines (n=33) culture-expanded from resected IBD patient tissue and stimulated for 3h with recombinant cytokines (adjusted P-values are shown where significantly different (p<0.05) compared to unstimulated, Kruskall-Wallis test). B) RNAseq analysis (Salmon log2-transformed TPM values, z-score, see STAR Methods) of cultured intestinal fibroblast cell line Ccd18-co, stimulated with either TNF-α (100ng/ml) or IL-1β (0.01ng.ml) for 3 hours (* P adjusted < 0.05 from DESeq2 differential gene expression analysis (see STAR Methods)). C) Dose-response of IL-1β and TNF-α stimulated Ccd18co fibroblasts for gene expression fold change (FC) of *CXLC8* over unstimulated, measured by qPCR. D) Pseudo-bulk expression fold changes (relative to M4/M5 low groups) of *ILR1* and *TNFR1* (see Supplementary Table 10) within the cellular clusters detected as in Figure 3A, across patients with either low, intermediate or high M4/M5 whole tissue expression. E) Gene set enrichment analysis of all modules detected in the discovery cohort for genes assigned to inflammasome pathways (GO:0061702).

**Supplementary Table 1. Clinical characteristics of the Oxford IBD patient discovery cohort used in this study**. Samples from the discovery cohort consist of surgically removed tissue of CD and UC patients (=IBD), as well as surgically removed normal tissue adjacent to colorectal tumours (= non-IBD). IBD, inflammatory bowel disease; CD, Crohn’s Disease; UC, Ulcerative colitis; IQR, interquartile range; n/a, not applicable.

**Supplementary Table 2. Detailed WGCNA analysis results**. Gene IDs and gene names of genes contained in the detected modules (modules_genes). Gene set enrichment analysis (fgsea) results of top 10 pathways upregulated in the detected modules (module GO analysis). Correlation strength (_Cor) and adjusted significance (_Pvalue) for correlation of individual modules with metadata traits across the inflamed IBD and CRC tissue samples (module-trait correlations).

**Supplementary Table 3. Replication of modules defined in the discovery cohort in other datasets**. Replication of the modules identified in the discovery cohort in publicly available datasets of IBD whole tissue gene expression.

**Supplementary Table 4. Differential expression of replication set modules in relation to therapy-response**. Significance test (Wilcoxon signed rank test) results for difference in module (eigengene value) expression between responders and non-responders to anti-TNF ^20^, corticosteroid ^15^ and anti-integrin ^21^ therapy, before the start of treatment.

**Supplementary Table 5. Predictive power of individual genes for therapy non-response**. Ranking of genes contained in all modules and detected in given dataset by area-under-the-curve (AUC) to predict non-response to anti-TNF ^20^, corticosteroid ^15^ and anti-integrin ^21^ therapy. Combined (summed) ranks for both anti-TNF and corticosteroid response are also shown.

**Supplementary Table 6. Clinical characteristics of the Oxford UC patient cohort of response to anti-TNF therapy**. Response to therapy in this UC patient cohort was defined as stopping anti-TNF therapy (Infliximab or Adalimumab) within 12 months of start, for reason of non-response (patients that stopped therapy for convenience, switch to biosimilar, or intolerance were not considered). Nancy histologic scores and UCEIS endoscopic scores, as well as the other characteristics, within 3 months before the start of anti-TNF therapy are shown. UC, Ulcerative colitis; IQR, interquartile range; UCEIS, Ulcerative Colitis Endoscopic Index of Severity.

**Supplementary Table 7. Clinical characteristics of the IBD patients used for RNAseq and FACS analysis**. Clinical characteristics of the IBD patient cohorts used for the transcriptomic and FACS analysis. UC, Ulcerative colitis; IQR, interquartile range; UCEIS, Ulcerative Colitis Endoscopic Index of Severity.

**Supplementary Table 8. Differential gene expression between neutrophils, stromal cells and mononuclear phagocytes FACS-sorted from IBD patient tissue**. List of all significant (adjusted P value < 0.05, |log2foldchange| >2) differentially expressed genes between neutrophils, stromal cells and MNPs sorted from the intestine of IBD patients. The standard DESeq2 outputs are reported.

**Supplementary Table 9. Differential gene expression between stromal cell clusters detected through scRNAseq**. Differential expression was performed to associate gene expression with stromal clusters comparing each cluster to all others irrespective of M4/M5 expression status. The standard DESeq2 outputs are reported.

**Supplementary Table 10. Differential gene expression between stromal cell clusters within M4/M5 high tissues only**. Differential expression was performed to associate gene expression with stromal clusters comparing each cluster to all others clusters within M4/M5 tissues only. The standard DESeq2 outputs are reported.

**Supplementary Table 11. Differential gene expression between inflammation states and clusters detected through scRNAseq**. For each gene (column feature), gene expression was associated with the tissue sample’s overall inflammatory status (column inflammatory_status), separately within each stromal cluster (column cell_type). The remaining columns are standard outputs of DESeq2.

**Supplementary Table 12. Differentially expressed genes in Ccd18-Co fibroblasts upon IL-1β stimulation**. List of all significant (adjusted P value < 0.05) differentially expressed genes in Ccd18-co fibroblasts after 3hour stimulations with IL-1β (0.01 ng/ml). The standard DESeq2 outputs are reported. log2FoldChange is the IL-1β-specific fold change over unstimulated condition.

