## Supplementary Table 6 for "IL-1-driven stromal-neutrophil interaction in deep ulcers defines a pathotype of therapy non-responsive inflammatory bowel disease"

| **Characteristic** | **UC responders to anti-TNF**  **(n=35)** | **UC non-responders to anti-TNF**  **(n=21)** |
| --- | --- | --- |
| % Male (Male/Female) | 65.7 (23/14) | 38.1 (8/13) |
| Median (IQR) age at sampling (years) | 32 (23-51) | 34 (24-52) |
| Median (IQR) disease duration at sampling (years) | 3 (1-9) | 2 (1-7) |
| UCEIS endoscopic score |  |  |
| Nancy histologic score (%)  = 0  = 1  = 2  = 3  = 4 | 0 (0)  1 (3.0)  8 (23.0)  18 (51.4)  10 (28.6) | 0 (0)  0 (0)  0 (0)  9 (42.9)  12 (57.1) |
| Medication (ever) during follow-up (%)  Aminosalicylates  Corticosteroids  Immunomodulator  (e.g. Azathioprine)  Treatment-naïve  (none of the above) | 26 (74.3)  18 (51.4)  17 (48.6)  0 (0) | 17 (81.0)  16 (76.2)  11 (52.4)  0 (0) |

**Supplementary Table 6. Clinical characteristics of the Oxford UC patient cohort of response to anti-TNF therapy.** Response to therapy in this UC patient cohort was defined as stopping anti-TNF therapy (Infliximab or Adalimumab) within 12 months of start, for reason of non-response (patients that stopped therapy for convenience, switch to biosimilar, or intolerance were not considered). Nancy histologic scores and UCEIS endoscopic scores, as well as the other characteristics, within 3 months before the start of anti-TNF therapy are shown. UC, Ulcerative colitis; IQR, interquartile range; UCEIS, Ulcerative Colitis Endoscopic Index of Severity.
