## Supplementary Table 7 for "IL-1-driven stromal-neutrophil interaction in deep ulcers defines a pathotype of therapy non-responsive inflammatory bowel disease"

| **Bulk RNAseq and FACS analysis (Figure 2)** | | | |
| --- | --- | --- | --- |
| **Characteristic** | **UC**  **N=9** | | **CD**  **N=3** |
| % Male (Male/Female) | 33.3(3/9) | | 100 (3/3) |
| Median (IQR) age at sampling (years) | 39 (20-72) | | 29 (22-50) |
| Median (IQR) disease duration at sampling (years) | 2 (0-25) | | 15 (5-34) |
| Nancy histologic score (%)  = 0  = 1  = 2  = 3  = 4 | 0 (0)  0 (0)  3(33.3)  4 (44.4)  2 (22.2) | | n/a |
| **single-cell RNAseq (Figures 3 and 4)** | | | |
| **Characteristic** | **UC=7** | **Healthy =4** | |
| % Female | 100% | 100% | |
| Median (IQR) age at sampling (years) | 56 (37-82) | 38 (21-73) | |
| Median (IQR) disease duration at sampling (years) | 17(4-62) | n/a | |
| Nancy histologic score (%)  = 0  = 1  = 2  = 3  = 4 | 1 (14.28)  0  2 (28.57)  4 (57.1) | n/a | |

**Supplementary Table 7. Clinical characteristics of the IBD patient cohorts used for RNAseq and FACS analysis.** UC, Ulcerative colitis; CD, Crohn’s Disease; IQR, interquartile range; n/a, not applicable.
