## Supplementary Table 1 for "IL-1-driven stromal-neutrophil interaction in deep ulcers defines a pathotype of therapy non-responsive inflammatory bowel disease"

| **Characteristic** | **non-IBD**  **(n=39)** | **IBD (UC+CD+IBDu)**  **(n=31)** | **UC**  **(n=8)** | **CD**  **(n=22)** |
| --- | --- | --- | --- | --- |
| % Male (Male/Female) | 54 (21/18) | 52 (16/15) | 60 (3/5) | 59 (13/9) |
| Median (IQR) age at sampling (years) | 68 (60-76) | 37 (29-49) | 36 (35-52) | 42 (29-54) |
| Median (IQR) disease duration at sampling (years) | n/a | 10 (5-20) | 5 (5-20) | 10 (5-20) |
| Sampling site (%)  Large intestine  Small intestine | 39 (100)  0 (0) | 23 (74)  8 (26) | 8 (100)  0 (0) | 14 (64)  8 (36) |
| Medication (ever) before surgery (%)  Aminosalicylates  Corticosteroids  Immunomodulator  (e.g. Azathioprine)  Anti-TNF  (Infliximab or Adalimumab)  Anti-Integrin  (Vedolizumab)  Monotherapy  (one of the above)  Combination therapy  (two or more of the above)  Treatment-naïve  (none of the above) | n/a  n/a  n/a  n/a  n/a  n/a  n/a  n/a | 6 (19)  12 (39)  21 (68)  13 (42)  2 (7)  10 (32)  15 (48)  6 (20) | 5 (63)  3 (38)  2 (25)  1 (6)  1 (6)  4 (50)  3 (38)  1 (12) | 1 (5)  9 (41)  11 (50)  7 (32)  1 (5)  6 (27)  12 (55)  4 (18) |

**Supplementary Table 1. Clinical characteristics of the Oxford IBD patient discovery cohort used in this study.** Samples from the discovery cohort consist of surgically removed tissue of UC, CD or IBDu patients (=IBD), as well as surgically removed normal tissue adjacent to colorectal tumors (= non-IBD). IBD, inflammatory bowel disease; CD, Crohn’s Disease; UC, Ulcerative colitis; IQR, interquartile range; n/a, not applicable.
